# CRISPR-repressed toxin-antitoxin provides population-level immunity against diverse anti-CRISPR elements

**DOI:** 10.1101/2024.01.05.574328

**Authors:** Xian Shu, Rui Wang, Zhihua Li, Qiong Xue, Jiajun Wang, Jingfang Liu, Feiyue Cheng, Chao Liu, Huiwei Zhao, Chunyi Hu, Jie Li, Songying Ouyang, Ming Li

## Abstract

Prokaryotic CRISPR-Cas systems are highly vulnerable to phage-encoded anti-CRISPR (Acr) factors. How CRISPR-Cas systems protect themselves remains unclear. Here, we uncovered a broad-spectrum anti-anti-CRISPR strategy involving a phage-derived toxic protein. Transcription of this toxin is normally reppressed by the CRISPR-Cas effector, but is activated to halt cell division when the effector is inhibited by any anti-CRISPR proteins or RNAs. We showed that this abortive infection-like effect efficiently expels Acr elements from bacterial population. Furthermore, we exploited this anti-anti-CRISPR mechanism to develop a screening method for specific Acr candidates for a CRISPR-Cas system, and successfully identified two distinct Acr proteins that enhance the binding of CRISPR effector to non-target DNA. Our data highlight the broad-spectrum role of CRISPR-repressed toxins in counteracting various types of Acr factors, which illuminates that the regulatory function of CRISPR-Cas confers host cells herd immunity against Acr-encoding genetic invaders, no matter they are CRISPR-targeted or not.

## INTRODUCTION

Bacteria utilize small CRISPR (clustered regularly interspaced short palindromic repeats) RNAs and Cas (CRISPR-associated) proteins to effectively combat various mobile genetic elements (MGEs), such as bacteriophages and^1–5^. The CRISPR-Cas systems, which exhibit a high level of diversity, can be categorized into two major classes, six types, and more than 30 subtypes^6^. When an MGE infects a bacterium, the CRISPR-Cas system selectively captures short fragments of foreign DNA, known as protospacers, which are flanked by a conserved protospacer adjacent motif (PAM)^7^. These captured protospacers are then integrated as spacers between repeated DNA segments within the CRISPR array, allowing the bacterium to effectively “memorize” information about the invader. The transcripts of CRISPR arrays are further processed to generate mature CRISPR RNAs (crRNAs). Each crRNA contains a specific spacer sequence that is flanked by repeat-derived handle sequences. Upon re-infection by MGEs, the CRISPR effector complex, comprising a crRNA and one or more Cas proteins (depending on the class), specifically targets and degrades the invading DNA, during which the recognition of PAM is essentially required to initiate target DNA binding^8,9^. In this manner, CRISPR-Cas systems equip bacterial cells with targeted immunity against MGEs.

In response, the rapidly evolving mobile genetic elements (MGE) have developed a variety of anti-CRISPR (Acr) proteins to counteract CRISPR immunity^10,11^. To date, more than 100 families of Acr proteins have been identified, with the majority known to inhibit type I CRISPR systems^12,13^. Most type I Acr proteins directly interact with the effector complex Cascade, sterically hindering the recognition of PAM, the binding of target DNA, or the recruitment of the helicase-nuclease Cas3 (responsible for cleaving the target DNA). Recent findings have further revealed some non-canonical inhibitory strategies, such as enzymatic modification of Cas proteins, induction of non-specific DNA binding, etc. (summarized in Ref. 13). Interestingly, a most recent study reported that bacteriophages can also utilize RNA anti-CRISPRs (Racrs) to suppress CRISPR immunity^14^. Racrs are encoded by solitary repeat units (SRUs) frequently observed in viral genomes^15^ and mimic crRNAs to interact with host Cas proteins, resulting in the formation of abnormal Cas subcomplexes^14^. Despite the remarkable diversity of viral anti-CRISPR strategies (either protein-based or RNA-based), the mechanisms by which CRISPR-Cas systems protect themselves from being blocked or suppressed remain poorly understood. In this study, we demonstrated that the recently unraveled CRISPR-repressed toxin-antitoxin (CreTA) modules^16^ can be activated by a wide range of diverse Acr proteins or Racr RNAs to trigger abortive infection, thereby expelling these Acr elements from the bacterial population and providing population-level protection for CRISPR-Cas.

CreTA was initially discovered by our group in the type I-B system of the archaeon *Haloarcula hispanica*^16^. The antitoxin, known as CreA RNA, is a crRNA analog that partially complements the promoter of the toxin. Based on this limited complementarity, CreA guides the CRISPR effector complex Cascade to suppress toxin expression at the transcription level, without incurring DNA cleavage. The toxin, CreT, is a small bacteriostatic RNA that inhibits cellular translation by sequestering a rare arginine transfer RNA (tRNA). Nevertheless, CreT RNAs from other species or CRISPR subtypes show poor sequence conservation, suggesting diverse mechanisms of toxicity^16^. For instance, CreT from the type I-B system of another archaeon *Halobacterium hubeiense* is a different bacteriostatic RNA that sequesters a rare isoleucine tRNA^17^. In contrast, CreT from the type I-F system of *Acinetobacter* sp.

WCHA45 acts as a bactericidal RNA, with its detailed mechanism yet to be elucidated^18^. Due to the lack of reported Acr proteins for these CRISPR systems, we first sought to explore additional CreTA modules from I-F systems in this study to investigate their potential anti-anti-CRISPR role (type I-F Acr proteins have been extensively discovered and characterized).

In the I-F CRISPR-Cas system of *Acinetobacter baylyi* ADP1, we identified a distinct CreTA module employing a phage-derived toxic protein, which we named CreP for ‘CRISPR-repressed proteinic toxin’. Our research showed that CreP is a bacteriostatic protein that inhibits cell division when it is over-expressed. Under physiological conditions, the promoter of CreP is repressed by CreA with the help of the CRISPR effector complex. Remarkably, we demonstrated that this CRISPR-repressed TA module can be triggered by a wide variety of MGE-encoded Acr proteins or RNAs that inactivate the CRISPR effector complex, thereby providing population-level resistance against these MGEs via abortive infection. Leveraging this broad-spectrum anti-anti-CRISPR activity, we also developed a screening strategy to identify Acr candidates capable of effectively suppressing a specific CRISPR-Cas system. Through this approach, we successfully identified two previously unknown Acr proteins, named AcrIF25 and AcrIF26. Interestingly, both of these proteins function by promoting non-specific DNA binding of the CRISPR effector complex. Altogether, these data highlight the broad-spectrum anti-anti-CRISPR role of CRISPR-repressed TA modules and their potential in conferring host cells the population-level immunity against Acr-encoding MGEs, which is independent of CRISPR targeting.

## Result

### A virus-derived toxin associates with *Acinetobacter* I-F CRISPR-Cas

By screening *Acinetobacter* genomes, we discovered a *creTA*-like module within the *cas3-csy1* intergenic region of the I-F CRISPR-Cas from *A. baylyi* ADP1 (Fig. 1a). The *creA* gene contains two CRISPR repeat-like sequences (referred to as ΨR1 and ΨR2, respectively) with a 30-bp spacing sequence (referred to as ΨS). Notably, the first 19 nucleotides of ΨS (excluding the 12^th^) exhibit a match to a target sequence preceded by a 5’-CC-3’ dinucleotide (Fig. 1a), which is a typical protospacer adjacent motif of the I-F CRISPR-Cas system^19^. Around 120 bp downstream of the CreA target, we observed a 324-bp open reading frame (ORF), which, interestingly, has been annotated as a ‘phage antirepressor KilAC domain-containing protein’. The KilAC domain was initially defined by Iyer *et al.* as ‘the C-terminal domain of phage P1 KilA’^20^. It was revealed that KilAC domain exhibits a highly mobile feature and is frequently fused with various N-terminal DNA-binding domains that are widespread in bacteriophages and eukaryotic DNA viruses^20^, such as the KilAN domain (N-terminal domain of phage P1 KilA) and Bro domain (N-terminal domain of baculovirus BRO proteins), as illustrated in Supplementary Fig. 1. Typically, these fused proteins are approximately 200-250 amino acids in length, while the ORF found between ADP1 *cas3* and *csy1* consists of only 103 amino acids, constituting a single KilAC domain. It is likely that this ORF originated from related phage proteins and was captured during the evolution of the *A. baylyi* ADP1 CRISPR-Cas locus.

**Fig. 1.**
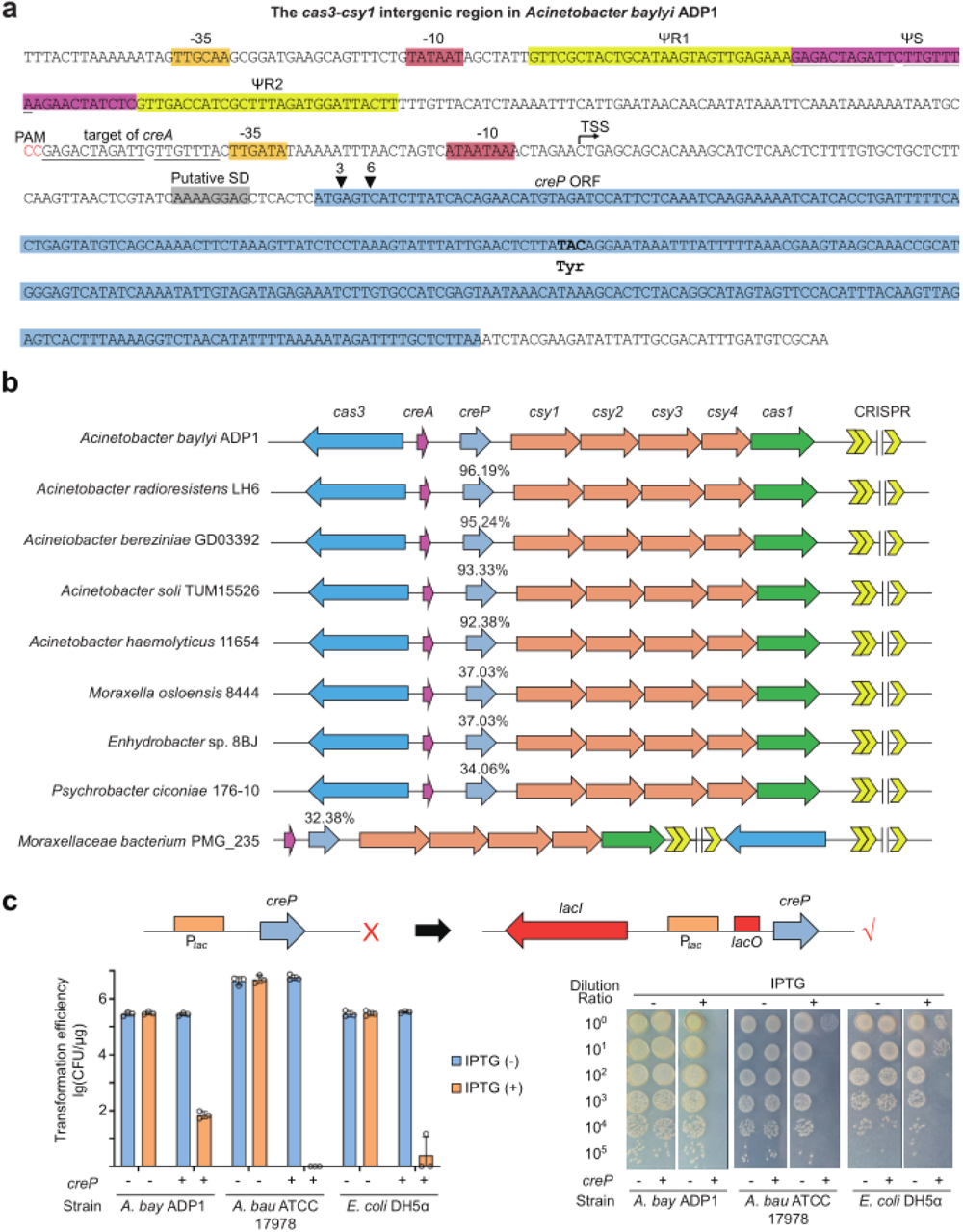
The putative CRISPR-repressed proteinic (CreP) toxin from *A. baylyi* ADP1. **a.** The *cas3-csy1* intergenic region (IGR) in *A. baylyi* ADP1 contains a *creAP* module. The *creA* gene has a ‘spacer’ sequence (ΨS) that is sandwiched by two CRISPR repeat-like sequences (ΨR1 and ΨR2). The predicted promoter elements (-10 and -35) of *creA* and *creP* are indicated. The transcription start site of *creP* was identified by primer extension (refer to Supplementary Fig. 4). The identical nucleotides between *creA* and the promoter of *creP* are underlined, with the protospacer adjacent motif (PAM) highlighted in red. The 43^rd^ codon TAC, encoding a tyrosine, was subjected to mutation (see Fig. 2). SD, Shine-Dalgarno sequence. ORF, open reading frame. **b.** Representative I-F CRISPR-Cas loci encoding CreP homologs. The sequence identity between ADP1 CreP and each homolog is indicated. **c,** Assessment of the toxic effect of ADP1 CreP. Placing *creP* under the control of a constitutive promoter (P*tac*) resulted in unsuccessful cloning in *E. coli* cells, while cloning with an IPTG-inducible P*tac* (containing *lacO*) proved to be feasible. Toxicity was initially assessed by transforming *A. baylyi* ADP1, *A. baumannii* ATCC 17978, and *E. coli* DH5α cells with plasmids bearing *creP*, and subsequently by dilution plating. The empty plasmid lacking *creP* served as a negative control (-). CFU, colony-forming units. Error bars, mean±s.d. (n=3).

Attempts to express this small ORF in *Escherichia coli* failed when using a constitutive *tac* promoter (P*_tac_*) (Fig. 1c), implying that its overexpression should be toxic, at least to *E. coli* cells. Thereby, we presumed that this toxic protein might be normally suppressed by CreA (and the Csy complex) in *A. baylyi* ADP1 cells, similar to the previously reported CRISPR-repressed RNA toxins within the CreTA modules^16,18^. To denote its putative proteinic nature, we hereinafter designated this KilAC domain-containing toxin as CreP for ‘CRISPR-repressed proteinic toxin’. Subsequently, we switched to an inducible P*_tac_* system (including the *lacO* operator, which is bound by the LacI repressor and responsive to IPTG induction), and achieved successful cloning (Fig. 1c). Then we observed that the CreP-expressing plasmid exhibited markedly reduced efficiency (compared to the empty vector) when transforming *A. baylyi* ADP1 (by ∼4 logs), *A. baumannii* ATCC 17978 (by ∼6.5 logs), or *E. coli* DH5α (by ∼5 logs) cells, and this reduction relied on IPTG induction (Fig. 1c). Therefore, CreP is toxic to a broad range of bacterial cells. In accordance, the dilution plating assay showed that CreP induction eradicated > 99.99% individuals of the *A. baylyi*, *A. baumannii*, or *E. coli* population (Fig. 1c).

By extensively searching the NCBI database, we obtained a total of 50 homologs of CreP, the encoding genes of which consistently reside within the *cas3-csy1* intergenic region of I-F CRISPR-Cas loci, with the exception of one case from *Moraxellaceae bacterium* PMG_235 (Fig. 1b and Supplementary Data. 1). It is worth noting that a significant majority, 84% to be precise (42 out of 50), of these *creP*+ CRISPR-Cas loci were from *Acinetobacter* genomes, while 10% (5 out of 50) were derived from *Moraxella osloensis* genomes (Supplementary Data. 1). These data illustrate the preference of CreP toxins for genomic association with I-F CRISPR-Cas systems, especially those from *Acinetobacter* species.

### CreP is a bacteriostatic protein

Thus far, three CreTA modules, respectively from the I-B CRISPR-Cas of *H. hispanica* ATCC 33960^16^, the I-B CRISPR-Cas of *H. hubeiense* JI20-1^17^, and the I-F CRISPR-Cas of *Acinetobacter* sp. WCHA45^18^, have been experimentally investigated. Intriguingly, although their *creT* genes each has a mini ORF, it has been demonstrated that the toxic element is not their protein but rather their RNA products. For instance, in the cases of *H. hispanica* and *H. hubeiense*, the CreT RNA sequesters the rare transfer RNA of arginine (tRNA^UCU^) and isoleucine (tRNA^CAU^), respectively, after translation initiation^16,17^. This information raises questions regarding the nature of the ADP1 CreP toxin, which we next aim to clarify.

We initially conducted the classic frameshift mutation analysis by inserting one, two, or three extra nucleotides after the first (base position 3) or the second trinucleotide (base position 6) of the CreP ORF (see Fig. 1a), and then introduced it into the native host *A. baylyi* ADP1 through transformation to assess its toxicity (Fig. 2a). Notably, we no longer observed a reduction in transformation efficiency caused by CreP induction when one or two nucleotides were inserted, which would have altered the reading frame. However, when three nucleotides were inserted (maintaining the reading frame), the toxicity of CreP was unaffected, and its induction reduced the transformation efficiency by a similar degree (approximately 5 logs) compared to the wild-type. Subsequently, we introduced a non-sense mutation to the CreP ORF by mutating the 43^rd^ codon TAC to the stop codon TAG, and then tested its toxicity in *E. coli* DH5α cells (Fig. 2b). As expected, this non-sense mutation deactivated the toxin, resulting in equivalent colony numbers when the cells were plated on either inducing or non-inducing medium. In contrast, expression of the wild-type CreP led to nearly complete eradication of *E. coli* cells, similar to the findings in Fig. 1c. Remarkably, when we *in trans* expressed a unique tRNA translating UAG to tyrosine (encoded by *supF* on a separate plasmid), the non-sense mutant of CreP regained its toxicity and its induction caused a comparable reduction in viable cells as observed with the wild-type CreP (Fig. 2b). These data substantially validate the proteinic nature of CreP.

**Fig. 2.**
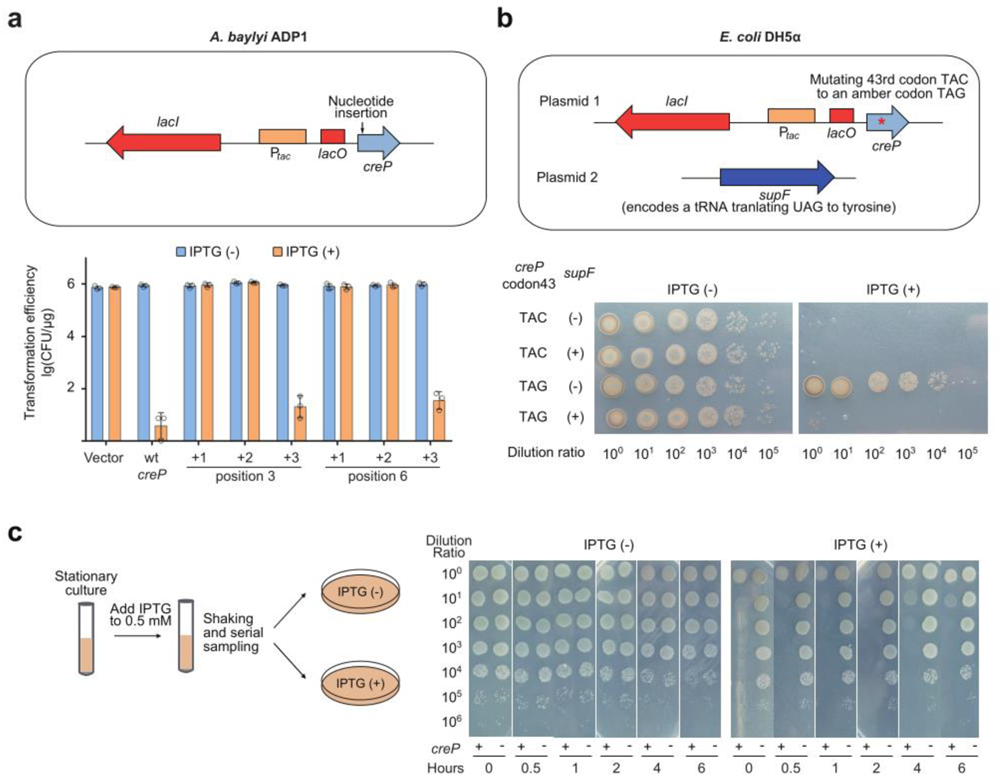
CreP is a bacteriostatic protein. **a.** Frameshift mutations were introduced into *creP* by inserting one (+1), two (+2), or three (+3) nucleotides immediately after the base position 3 or 6 (indicated in Fig. 1a). The toxicity was evaluated by transforming *A. baylyi* ADP1 cells with plasmids expressing the wild-type (wt) or mutated *creP*. Error bars, mean±s.d. (n=3). **b.** Non-sense mutation (changing a tyrosine codon TAC to the stop codon TAG) of *creP* and its suppression by a tRNA translating UAG to tyrosine (encoded by *supF*). The toxicity was evaluated by plating *E. coli* DH5α cells containing the wild-type (codon43: TAC) or mutated (TAG) *creP* on medium with or without IPTG. **c,** Dilution plating of *A. baylyi* ADP1 cells (containing the IPTG-inducible *creP*) at 0.5, 1, 2, 4, or 6 hours after IPTG induction.

Next, we asked whether CreP functions as a bactericidal or bacteriostatic toxin. We inoculated the *A. baylyi* ADP1 cells containing an inducible *creP* gene into non-inducing medium (alongside a control group containing the empty vector), and only after the culture reached stationary phase (OD approaching 1), was CreP expression induced (Fig. 2c). At various time intervals post induction, the cell culture was sampled, serially diluted, and then plated onto both inducing and non-inducing medium simultaneously. As expected, the *creP*+ ADP1 cells hardly grew on the inducing medium. Interestingly, on the non-inducing medium, the *creP*+ and *creP*-cells displayed similar colony forming unit (CFU) counts at each specific time point (0.5, 1, 2,4, or 6 hours post induction) (Fig. 2c), indicating continuous CreP expression up to 6 hours did not reduce the number of viable cells in the stationary culture. This data leads us to conclude that CreP is a bacteriostatic protein.

### CreP inhibits cell division

The KilAC domain can exist independently (like in CreP homologs), and as mentioned above, it can also fuse with the KilAN domain (like in the KilA protein of the *Escherichia coli* phage P1^21^) or with the Bro domain (like in the gp07 of the *Staphylococcus aureus* phage Phi11^22^). By multiple sequence alignment, we observed that CreP homologs are highly conserved in primary sequence and share limited conserved residues with the carboxyl-terminal of P1 KilA protein, such as the ‘AKxLxL’ and ‘TxKG’ combinations (Fig. 3a). Importantly, the predicted three-dimensional structure of ADP1 CreP exhibits a similar architecture to the carboxyl-terminal of P1 KilA or Phi11 gp07 (Fig. 3b). It has been reported that the P1 KilA and Phi11 gp07 proteins not only act as anti-repressors during the lytic development of their respective phages, but also have the ability to inhibit host cell division^22,23^. Specially, it has been shown that the heterologous expression of Phi11 gp07 in *E. coli* leads to cell filamentation^22^. Therefore, we examined the changes in morphology of *A. baylyi* ADP1 cells after inducing the expression of CreP. Remarkably, the majority of cells expressing CreP formed very long filaments (mostly measuring 20-30 μm), while the control cells (containing an empty vector) maintained their regular cell size (2-3 μm) (Fig. 3c). When *E. coli* cells expressing CreP proteins were examined under the microscope, similar cell filamentation was observed (Supplementary Fig. 2). Thus, similar to the distantly related phage antirepressors, CreP inhibits cell division, leading to cell filamentation.

**Fig. 3.**
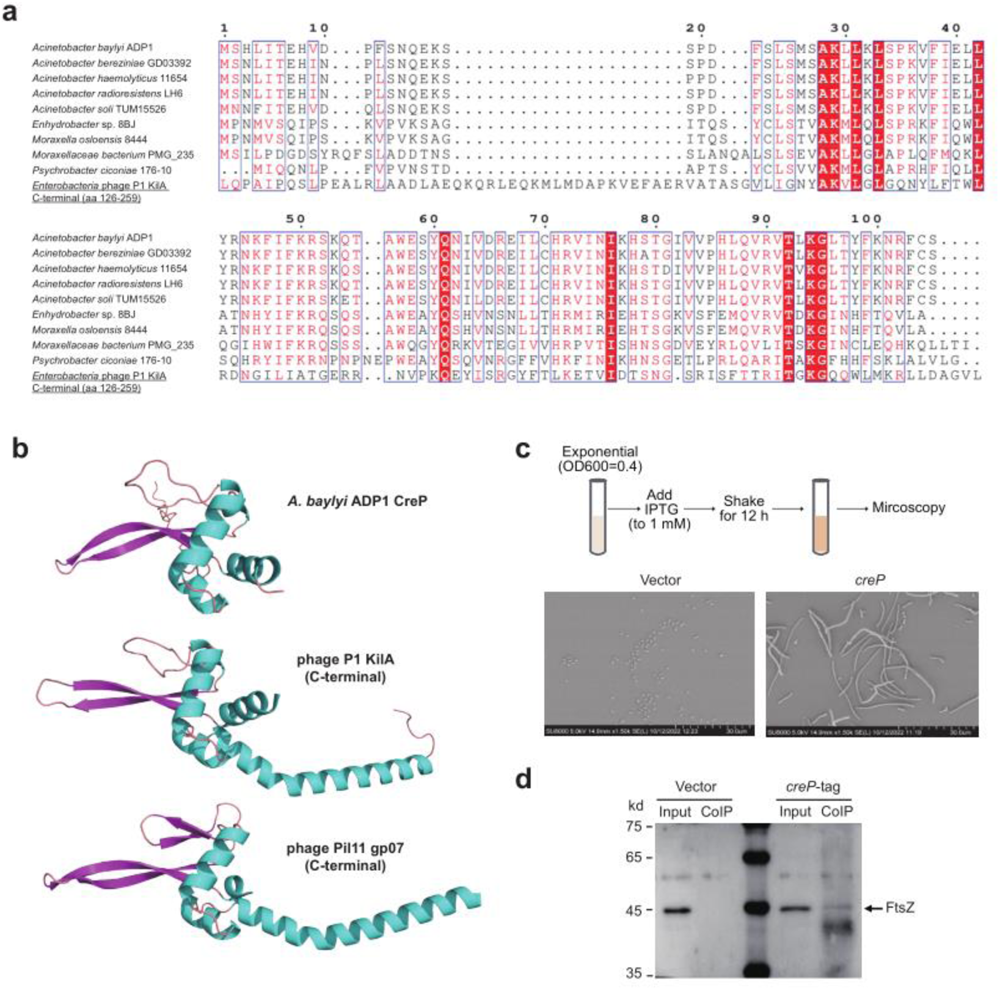
CreP overexpression induces cell filamentation. **a.** Multiple sequence alignment of CreP and its homologs. The C-terminal KilA domain of a phage antirepressor encoded by the *Enterobacteria* phage P1 was included for alignment. Numbers indicate the amino acid position in ADP1 CreP. **b.** Predicted structure of ADP1 CreP and its homology to the C-terminal of phage P1 KilA and phage Pih11 gp07 proteins. **c.** Microscopy of *A. baylyi* ADP1 cells over-expressing CreP or containing an empty vector. **d.** Co-immunoprecipitation (CoIP) assay followed by Western blot to examine the interaction between CreP and FtsZ. Flag-His6-tagged CreP was used as the bait in the CoIP assay and an empty vector was included as a mock control. Input represents the cellular extract used for CoIP assays.

In most bacteria, cytokinesis involves the assembly of FtsZ, a highly conserved prokaryotic tubulin-homolog, into a ‘Z-ring’ structure at the site of cell division^24^. We expressed a tagged CreP protein in *E. coli* cells, and then used it as a bait in a co-immunoprecipitation assay to collect cellular proteins that interact with CreP. Subsequently, we conducted a Western blot using an antiserum against purified FtsZ and observed a weak corresponding band among the co-immunoprecipitated proteins (Fig. 3d). This suggests that CreP has the ability to interact with FtsZ, either directly or indirectly. However, further investigation is required to determine whether there are additional cellular targets for CreP.

### The native promoter of *creP* is repressed by CreA RNA

The finding that overexpression of CreP in its native host *A. baylyi* ADP1 resulted in cell filamentation indicates that the natural *creP* gene on the ADP1 chromosome is tightly regulated under physiological conditions. Our recent studies have revealed that different CreT RNA toxins from *H. hispanica*, *H. hubeiense*, and *Acinetobacter* sp. WCHA45, are all transcriptionally repressed by their corresponding CreA RNAs with the help of the CRISPR effector complex^16–18^. Based on this information, we hypothesize that in *A. baylyi* ADP1, the predicted *creA* gene located upstream of *creP* (see Fig. 1a) may also produce a guide RNA that directs the Csy complex to inhibit *creP* transcription.

We first analyzed the ‘repeat’ elements (ΨR1 and ΨR2) of *creA*. Despite containing 9 and 16 differing nucleotides, respectively, compared to the conserved CRISPR repeat, both elements maintained a pair of internal inverted repeats, indicating their potential to form a stem-loop RNA structure that can be recognized by the endonuclease Csy4 (Fig. 4a). Nevertheless, it is worth noting that the inverted repeats within ΨR2 are separated by one additional nucleotide compared to those of CRISPR repeat and ΨR1. Through small RNA sequencing, we were able to determine the precise sequence of mature CreA RNA, which surprisingly consists of the last 8 nucleotides of ΨR1 (a typical 5’ handle derived from Csy4 processing) and only the first 25 nucleotides of ΨS (Fig. 4b and Supplementary Fig. 3). We speculate that the ΨS sequence may contain a transcription terminating site or a processing site of unknown nuclease(s).

**Fig. 4.**
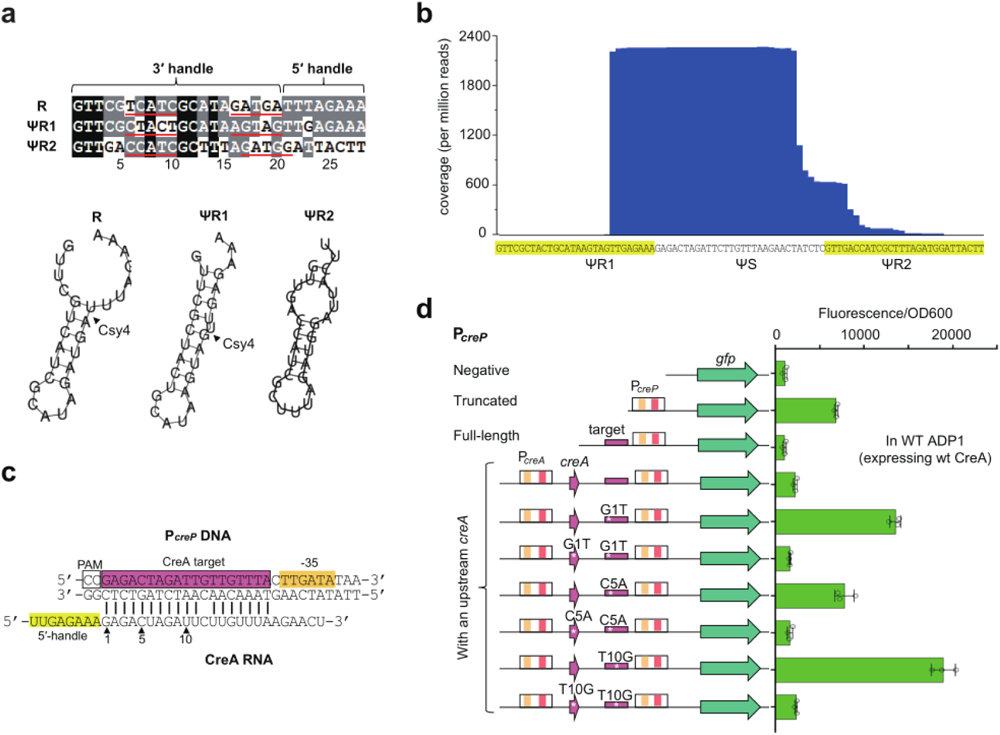
Characterization of CreA sequence and its inhibition of P*creP*. **a.** Homology between the CRISPR repeat (R) and the *creA* ‘repeats’ (ΨR1 and ΨR2) of *A. baylyi* ADP1. Nucleotides corresponding to the 5’ or 3’ handle on mature RNA molecules are indicated. Red lines indicate palindromic nucleotides with the potential to form a stem on the RNA secondary structure (where the cleavage site of Csy4 is indicated). **b.** Determination of the sequence of mature CreA RNA through small RNA-seq. **c.** Complementarity between CreA RNA and the promoter DNA of *creP.* The promoter element -35 is shown. The 1^st^, 5^th^, and 10^th^ base pairings relative to PAM (protospacer adjacent motif) were subjected to separate complementary mutation analysis (see panel **d**). **d.** Repression effect of CreA on a P*creP*-controlled *gfp* gene. White asterisks indicate point mutations (G1T, C5A, and T10G) to disrupt or restore the complementarity between the plasmid-born *creA* and P*creP*. Note that this assay was performed in wild-type ADP1 cells that contained a chromosomal copy of *creAP* (wild-type). Error bars, mean±s.d. (n=3).

In Fig. 4c, we illustrate the complementarity between the mature CreA RNA and its putative target within the promoter of *creP* (P*_creP_*). Through primer extension, we identified the transcription start site (TSS) of *creP* (Supplementary Fig. 4) and accordingly predicted the -10 and -35 elements that are crucial for promoting transcription (Fig. 1a). Note that the -35 element is located immediately downstream of the target site of CreA (Fig. 4c), implying that the binding of the Csy-CreA ribonucleoprotein complex to P*_creP_* DNA may sterically block the initiation of transcription. Utilizing the *gfp* (green fluorescence protein) reporter gene, we showed that a truncated P*_creP_* lacking the CreA target site showed approximately 6.6 times higher activity than the full-length P*_creP_* in ADP1 cells expressing the wild-type CreA (Fig. 4d), which supports the inhibitory effect of CreA on P*_creP_*. Then we cloned a 331-bp fragment containing *creA* (including its predicted promoter), the target sequence of *creA*, and P*_creT_* to control the expression of *gfp* (Fig. 4d). Interestingly, fluorescence driven by this fragment was ∼2.2 times higher than that driven by P*_creP_* alone, indicating the influence of upstream sequences on P*_creP_*. Based on this design, we separately mutated the 1^st^, 5^th^, and 10^th^ nucleotides (with respect to PAM) of the CreA target sequence to disrupt the CreA-P*_creP_* complementarity, which increased fluorescence by a factor of 6.0, 3.5, and 8.4, respectively (Fig. 4d). Notably, when the upstream *creA* gene was correspondingly mutated to restore the complementarity, fluorescence declined to the level observed in the wild-type design. These data provide substantial evidence that CreA represses P*_creP_* based on sequence complementarity.

### CreP can be activated by a wide variety of Acr proteins

Next, we investigated the physiological role of CreP during CRISPR immunity. Because CreP expression is normally repressed by CreA and the Csy complex, we hypothesized that phage-encoded Acr proteins which inhibit the Csy complex could potentially trigger the toxicity of CrePA. Although numerous Acr proteins have been identified for the *Pseudomonas aeruginosa* I-F CRISPR-Cas systems, none have been reported for any *Acinetobacter* CRISPR-Cas systems. Therefore, we initially synthesized a range of *P. aeruginosa acrIF* genes (*acrIF1*-*acrIF14*) and introduced them into *A. baylyi* ADP1 cells to investigate their ability to inhibit the ADP1 CRISPR-Cas and induce CreP toxicity (Supplementary Fig. 5a). However, all *acrIF*-expressing plasmids exhibited a transform efficiency comparable to the empty vector, with the exception of *acrIF5*, which caused only a mild reduction in transformation efficiency (by about 2 logs) (Supplementary Fig. 5b). By transforming a Δ*crePA* mutant, we demonstrated that this reduction was independent of CrePA (Supplementary Fig. 5c), which suggests that AcrIF5 *per se* is toxic to *A. baylyi* ADP1 cells. Therefore, it appears that the *P. aeruginosa* AcrIF proteins display a high degree of specificity and are unable to inhibit the *A. baylyi* I-F CRISPR-Cas system.

To circumvent this issue, we modified the *crePA* module by replacing ΨR1 and ΨR2 with the conserved repeat of the *P. aeruginosa* PA14 CRISPR array to make it fitting with the *P. aeruginosa* Csy complex (Fig. 5a), because our recent study showed that the specificity between CreTA and CRISPR-Cas can be readily altered through ‘repeat replacement’^25^. We denoted the modified *creA* gene as *creA\**. Next, we introduced a plasmid carrying the PA14 *csy* operon and the *crePA\** module into *A. baumannii* ATCC 17978, which lacks a native CRISPR-Cas system and is susceptible to CreP toxicity (Fig. 1c). Subsequently, we transformed the modified *A. baumannii* ATCC 17978 cells with plasmids expressing individual *P. aeruginosa* AcrIF proteins. Strikingly, expression of most AcrIF proteins markedly reduced transformation efficiency (by 4 to 5 logs), except for AcrIF3, AcrIF4, and AcrIF5 (Fig. 5b). Interestingly, while most AcrIF proteins disrupt the specific binding of the Csy complex to target DNA in various ways^12,13^, AcrIF3 and AcrIF5 were found to inhibit the recruitment of Cas3 helicase-nuclease to the Csy-DNA compelx^10,26^. It is worth noting that AcrIF4 expression still caused a slight decrease in transformation efficiency (∼50%; *P* = 0.0059) and resulted in smaller colonies on selective plates (Fig. 5c), which coincides with an *in vitro* study that showed AcrIF4 only reduces, but does not completely abolish, the binding between Csy and target DNA^27^. Additionally, the mechanism of AcrIF12 remains unknown, but its expression was found to induce the toxicity of CrePA (Fig. 5b), suggesting that it possibly, like most AcrIF proteins, disrupts the target DNA binding activity of the Csy complex.

**Fig. 5.**
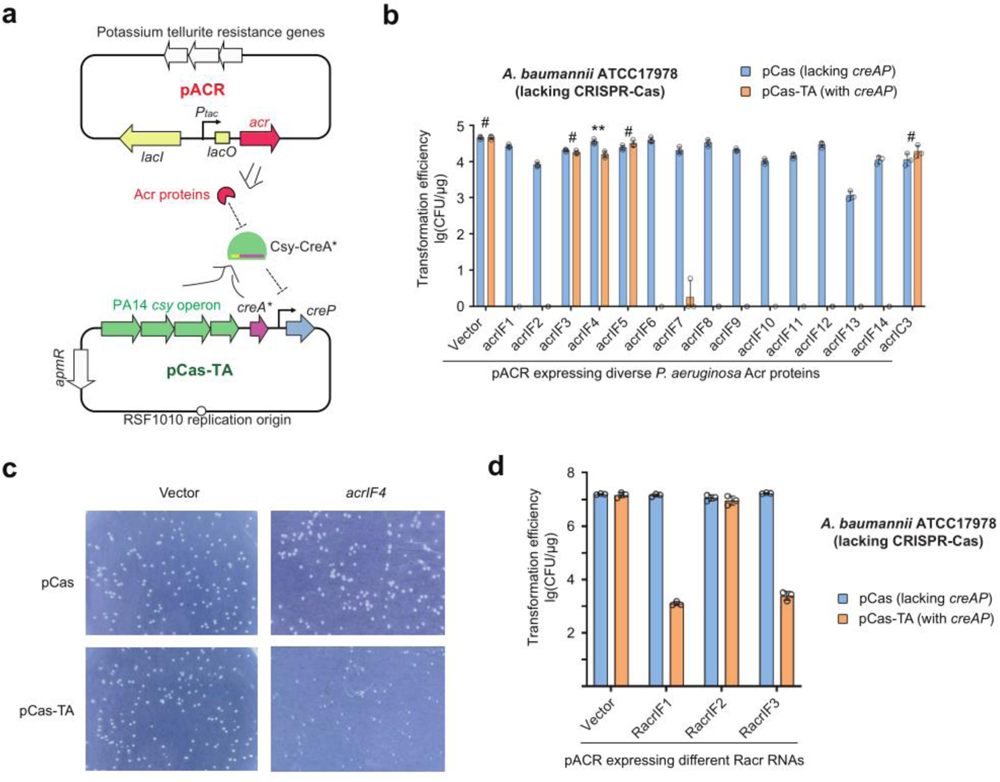
CreP toxicity can be triggered by various Acr proteins and Racr RNAs. **a.** Illustration of the activation of *creP* by *P. aeruginosa* AcrIF proteins. The ADP1 *creAP* module was modified (ΨR1 and ΨR2 of *creA* were replaced by the CRISPR repeat of *P. aeruginosa* PA14) to fit with PA14 Csy complex, and the modified *creA* is referred to as *creA\**. pCas-TA carries the PA14 *csy* operon, *creA*\*, and *creP*. pACR carries a *P. aeruginosa acrIF* gene controlled by the IPTG-inducible P*tac*. **b.** Transformation of *A. baumannii* ATCC17978 cells (containing pCas-TA or the *crePA*-lacking pCas) with different pACR plasmids. *acrIC3* served as a negative control. Error bars, mean±s.d. (n=3). # *P*>0.05. \*\**P*<0.01. **c.** Transformant colonies formed on selective plates. **d.** Transformation of *A. baumannii* ATCC17978 cells (containing pCas-TA or the *crePA*-lacking pCas) with plasmids expressing different Racr RNAs. Error bars, mean±s.d. (n=3).

A very recent study revealed that bacteriophages can also suppress CRISPR-Cas by employing RNA-based anti-CRISPRs (Racrs), which specifically interact with Csy4 and Csy3 to form aberrant Cas subcomplexes^14^. This study also identified three Racrs, namely RacrIF1-3, that have the ability to inhibit the CRISPR-Cas system of *P. aeruginosa* PA14. To investigate their impact on CrePA, we transformed *A. baumannii* ATCC 17978 cells (containing pCas-TA or pCas) with Racr-expressing plasmids. As expected, expression of RacrIF1 or RacrIF3 markedly reduced transformation efficiency (by approximately 4 logs) compared to the empty vector, and this effect relied on the presence of CrePA (Fig. 5d). However, the same effect was not observed for RacrIF2, for reasons that are currently unknown. In conclusion, CrePA is a TA-like module capable of reacting to various Acr or Racr elements that influence the target DNA binding activity or the normal assembly of the CRISPR effector. Conceivably, such CRISPR-repressed TA modules may provide CRISPR-Cas with population-level resistance against diverse Acr proteins via inducing abortive infection (see below).

### The CrePA module can be harnessed to screen new Acr proteins

We wondered whether the broad-spectrum anti-Acr role of CrePA could be exploited to screen for new Acr proteins that inhibit a specific CRISPR-Cas system, e.g., the one of *A. baylyi* ADP1 (Fig. 6a). We first searched the *Acinetobacter* genomes for homologs of the known *P. aeruginosa* Acr proteins, and preliminarily identified four *Acinetobacter* homologs for AcrIF6 (designated AcrIF6.2-AcrIF6.5) and for AcrIF11 (designated AcrIF11.2-AcrIF11.5), respectively. We then examined their anti-CRISPR activity against the ADP1 CRISPR-Cas by simply transforming the ADP1 cells with plasmids expressing one of these predicted Acr proteins. Remarkably, the induced expression of AcrIF6.2, AcrIF11.3, and AcrIF11.5 significantly reduced the transformation efficiency (by approximately 6 logs) (Fig. 6b). To rule out the possibility that these Acr proteins were toxic to ADP1 cells themselves (as observed for AcrIF5; Supplementary Fig. 5), we separately introduced their expression plasmids into wild-type and Δ*crePA* ADP1 cells. Notably, these three Acr proteins only reduced the transformation efficiency of wild-type cells (containing *crePA*), but not that of Δ*crePA* ADP1 cells (Fig. 6c), illustrating that these proteins are not toxic *per se* and should have induced the toxicity of CrePA.

**Fig. 6.**
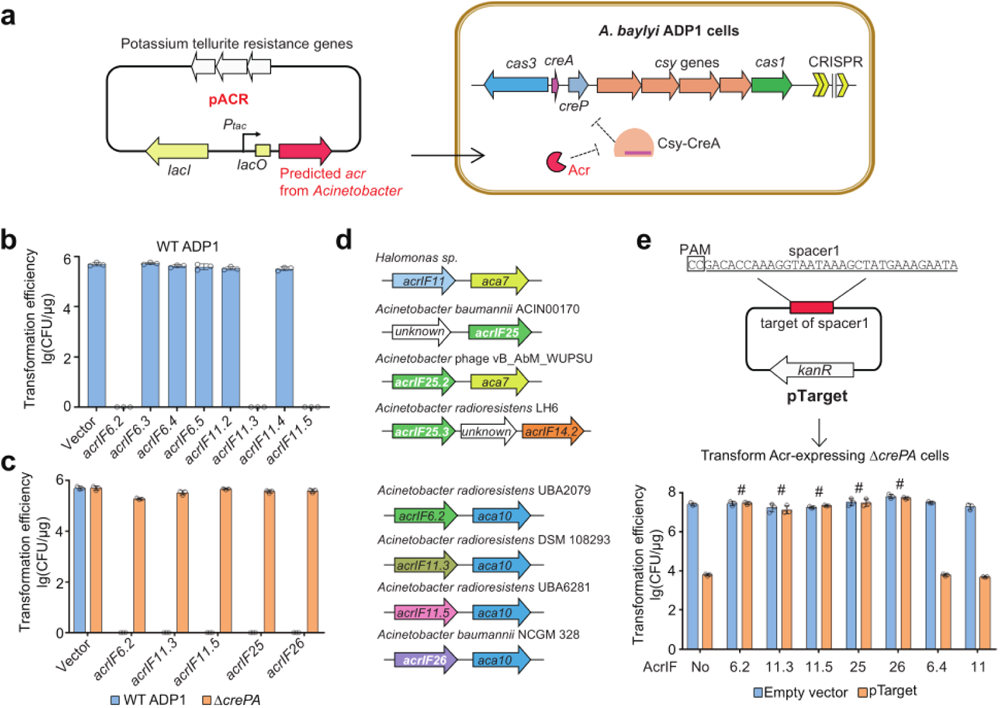
CrePA can be utilized for screening new Acr candidates. **a.** Schematic representation of the CrePA-based assay for screening new Acr proteins that inhibit *A. baylyi* ADP1 CRISPR-Cas. Inhibition of CRISPR-Cas is expected to activate the CRISPR-repressed *creP*, leading to cellular toxicity. **b.** Transformation efficiency of wild-type (WT) ADP1 cells by pACR derivates, which carries the *Acinetobacter* homologs of *acrIF6* or *acrIF11*. Error bars, mean ± s.d. (n=3). **c.** Transformation of ADP1 cells with plasmids containing newly predicted *acrIF* genes. Δ*crePA*, a mutant lacking *creAP*. Error bars, mean±s.d. (n=3). **d.** Prediction of new *A. baumannii acrIF* genes based on their genomic association with *aca7* or *aca10*. **e.** Validation of the anti-CRISPR function of explored *A. baumannii* Acr proteins. Δ*crePA* cells with plasmid-expressed Acr proteins were transformed by pTarget, which is targeted by the first spacer (spacer1) of the ADP1 CRISPR array. AcrIF6.4 and AcrIF11, which cannot inhibit ADP1 CRISPR-Cas, served as negative controls. PAM, protospacer adjacent motif. Error bars, mean±s.d. (n=3).

In addition to the sequence similarity search, we also searched for new Acr proteins based on their genetic link with Acr-associated (*aca*) genes, which encode helix-turn-helix (HTH) proteins that regulate Acr expression. Using the *aca7* gene associated with *acrIF11* and a new *aca* gene (designated *aca10*) associated with *acrIF6.2*, *acrIF11.3*, and *acrIF11.5*, we identified two novel candidate *acr* genes from the *Acinetobacter* genomes, designated *acrIF25* and *acrIF26*, respectively (Fig. 6d). By introducing these two genes into ADP1 cells, we demonstrated that expression of either gene caused a marked reduction in transformation efficiency, which notably, depended on the presence of *crePA* (Fig. 6c). Therefore, the newly identified AcrIF25 and AcrIF26 can both trigger the toxicity of CrePA, conceivably by inhibiting the ADP1 CRISPR-Cas. Finally, we designed a target plasmid bearing the protospacer (with PAM) of the first spacer in the ADP1 CRISPR array, and used it to transform Δ*crePA* cells expressing a specific Acr protein (Fig. 6e). In cells that did not express any Acr proteins or those expressing a *P. aeruginosa* Acr protein (Acr6 or Acr11) that did not induce CrePA toxicity (refer to Supplementary Fig. 5), CRISPR immunity against this target plasmid was not influenced and resulted in a significant 3-log reduction in transformation efficiency (Fig. 6e). In contrast, this plasmid interference effect was not observed in cells that express any of the Acr proteins capable of inducing the toxicity of CrePA (Fig. 6e). This data confirms that the candidates observed to induce the CrePA toxicity of *A. baylyi* ADP1 are *bona fide* Acr proteins that actively inhibit the ADP1 CRISPR-Cas. By introducing these Acr proteins into ADP1 cells containing a P*_creP_*-controlled *gfp* gene, we further demonstrated their efficacy in relieving the suppression of P*_creP_* (Supplementary Fig. 6). This confirms that these Acr proteins function by inhibiting the target DNA binding of the Csy complex. Notably, each Acr exhibited a distinct relieving effect on P*_creP_* repression, with fluorescence increasing by a factor ranging from 3.5 to 11.8 compared to the Acr-control (Supplementary Fig. 6), implying their different mechanisms of inhibiting the Csy complex.

### Both AcrIF25 and AcrIF26 induce the non-specific DNA binding of Csy complex

Because AcrIF25 and AcrIF26 lack sequence similarity to any previously reported Acr proteins, we sought to investigate their anti-CRISPR mechanisms. By size-exclusion chromatography, we showed that both AcrIF25 and AcrIF26 can directly bind to the purified *A. baylyi* ADP1 Csy complex (Fig. 7a and 7b). Then we performed EMSA to examine whether these proteins could inhibit the target DNA binding of the Csy complex *in vitro*. Remarkably, we observed no inhibitory effects for AcrIF25, event at a concentration 9 times higher than the Csy complex (Fig. 7c), and interestingly, at high concentrations, AcrIF26 slightly enhanced the target DNA binding (Fig. 7d). We then repeated this assay using a non-target DNA substrate and found that both Acr proteins effectively promote the binding of Csy to the non-target DNA (Fig. 7d). Intriguingly, at a high concentration, AcrIF26 appeared to induce the dimerization of the Csy-Acr-DNA ternary complex.

**Fig. 7.**
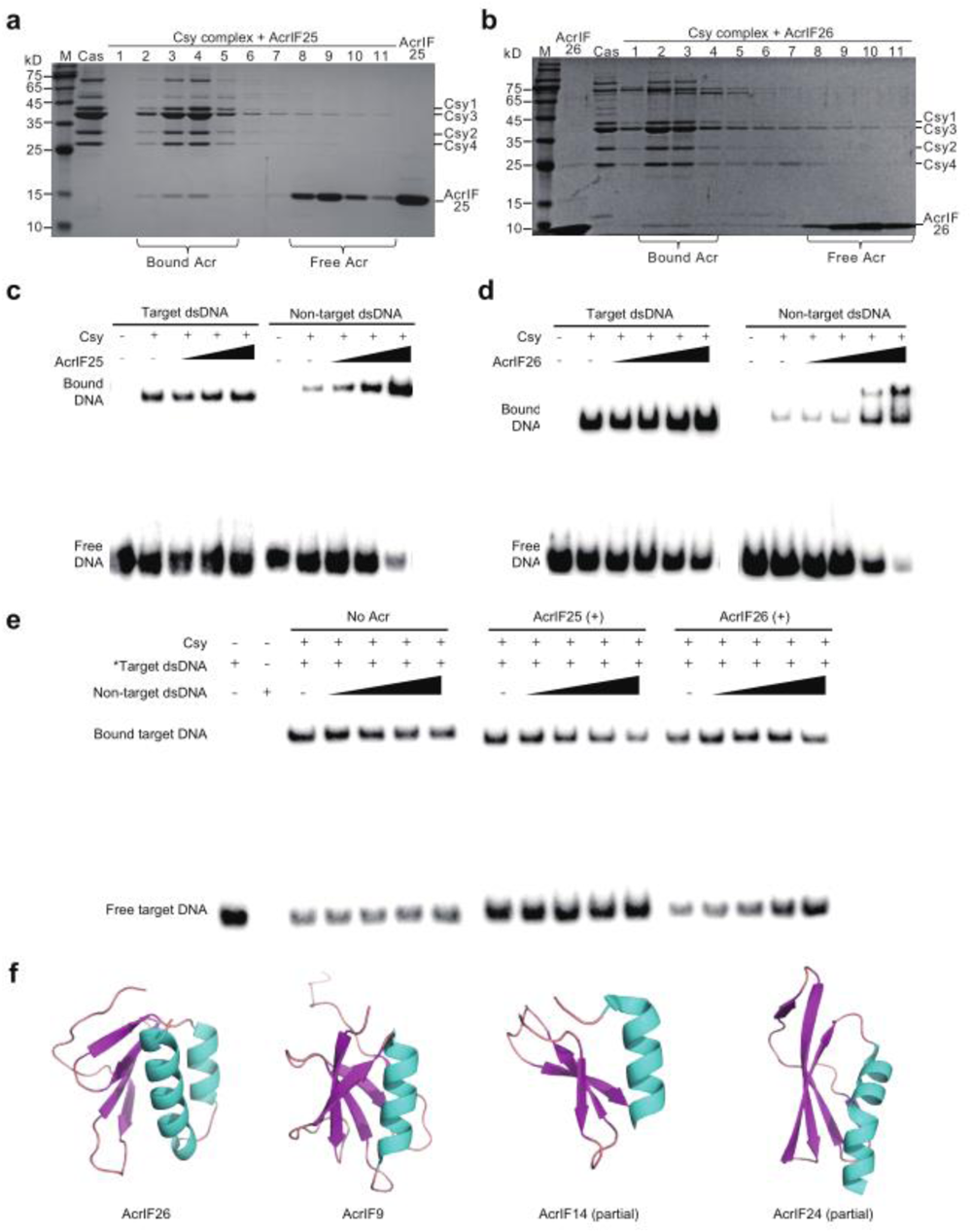
Both AcrIF25 and AcrIF26 induce the binding of Csy complex to non-target DNA. **a** and **b.** Examine the direct interaction between ADP1 Csy complex and AcrIF25/AcrIF26 by size-exclusion chromatography (SEC). Purified AcrIF25 (**a**) or AcrIF26 (**b**) was incubated with purified Csy complex and the mixture was fractionated by SEC. The SEC fractions were analyzed by SDS-PAGE gel in numbers according to their elution position. The purified Acr and Csy complex are included as controls. Protein bands are indicated by arrows. M, protein ladders in kD. **c** and **d.** EMSA assays to examine the effect of AcrIF25 (0, 9, 27, or 81 nM) or AcrIF26 (0, 9, 27, 81, or 243 nM) on the binding affinity of target DNA or non-target DNA to Csy complex (9 nM). **e.** Competition of non-target DNA (0, 2, 4, 8, or 16 nM) with biotin-labeled target DNA (0.5 nM) for 9 nM Csy complex in the absence or presence of Acr proteins (240 nM). **f.** Predicted structure of AcrIF26 and its structural homology to reported AcrIF proteins that also induce the non-specific DNA binding of I-F Csy complex.

To investigate whether the binding sites of target and non-target DNA overlap, we further conducted the competition EMSA experiments. Biotin-labeled target DNA was pre-mixed with Csy in the presence or absence of AcrIF25/AcrIF26, and then unlabeled non-target DNA was added in increasing concentrations. In the absence of Acr proteins, the ratio of free target DNA remained largely unaffected by the addition of non-target DNA. However, when AcrIF25 or AcrIF26 was present in the competition assay, the ratio of free target DNA increased as the concentration of non-target DNA increased (Fig. 7d). Therefore, AcrIF25 and AcrIF26 promote non-target DNA to compete with target DNA for the same or overlapping binding site on the Csy complex.

We predicted the structure of AcrIF25 and AcrIF26 by AlphaFold2, and conducted a search in the PDB database based on structure similarity. We found that AcrIF26 is composed of a four-stranded anti-parallel β sheet and two helices and shows a significant resemblance to *P. aeruginosa* AcrIF9 (Fig. 7f), which also promotes the binding of non-target DNA by Csy. We also noted that these two Acr proteins share a similar arrangement of secondary structures with the C-terminal domain of AcrIF14 and AcrIF24, both of which also induce the non-target DNA binding of Csy (Fig. 7f). This suggests that AcrIF26 employs a relatively conserved structure among the different Acr proteins that induce non-specific DNA binding of the Csy complex.

Notably, AcrIF25 was predicted to have a distinct two-domain structure, comprising an N-terminal helix-turn-helix (HTH) domain (residues 1–79, NTD) and a C-terminal β barrel domain (residues 1–79, CTD) consisting of four twisted β strands (Supplementary Fig. 7a and 7b). The presence of the HTH domain suggests the potential of AcrIF25 to act as an Aca protein, reminiscent of the Acr-Aca protein AcrIF24^28,29^. We attempted to determine the crystal structure of AcrIF25, but possibly due to the high flexibility of the loop connecting CTD and NTD, we were only successful in solving the structure of the HTH NTD (Supplementary Data. 2; PDB code: 8X24). The AcrIF25 NTD forms a homodimer through extensive hydrophobic interactions and hydrogen bonding observed between residues at the dimer interface (Supplementary Fig. 7c and 7d). For example, residues Ile52, Leu55, Leu59, Ile65, Leu66, Leu69, Leu70 and Leu73 in α4 of molecule A interacted with their equivalent residues in α4’ of molecule B, forming a hydrophobic core (Supplementary Fig. 7e). Additionally, residues Arg62 and Leu73 in α4 formed hydrogen bonds with residues Leu8 and Ala9 in α1’ and with residue Lys56 in α4’, respectively, reinforcing the dimer formation (Supplementary Fig. 7f). Through a cross-linking experiment, we confirmed that AcrIF25 primarily exists as dimers (Supplementary Fig. 7g), which suggests its potential to interact with Csy complexes in a dimer state.

### CrePA expels Acr-encoding mobile elements from the bacterial population

At last, we evaluated the effect of CrePA in protecting CRISPR-Cas against Acr proteins in a bacterial population. We constructed two plasmids: one carries the reporter gene *gfp* and no *acr* genes (pGFP), while the other contained the reporter gene *mcherry* and an IPTG-inducible *acr* gene (pMCherry) (Fig. 8a). These plasmids were mixed in equal amounts and introduced into wild-type *A. baylyi* ADP1 cells through electro-transformation. The transformed cells were co-cultured and sub-inoculated each time the culture reached the stationary phase. Note that IPTG was added to the culture during the second and third sub-inoculations to induce Acr expression. For each stationary-phase culture, we measured the green and red fluorescence simultaneously (Fig. 8a) and examined the cells using fluorescence microscopy (Fig. 8b). Notably, when pMCherry contained the newly identified AcrIF25 or AcrIF26, which inhibit the ADP1 CRISPR-Cas system, the red fluorescence rapidly decreased and eventually almost disappeared 20 hours after Acr induction (Fig. 8a). In parallel, the green fluorescence (produced by pGFP that did not encode Acr proteins) increased accordingly. This indicates that cells containing the Acr-expressing plasmid were extinct from the population, as confirmed by the complete absence of red cells in the microscope field of view (Fig. 8b). It should be noted that when pMCherry encoded AcrIF6.4, which was shown to be unable to inhibit the CRISPR-Cas of ADP1 cells, the extinction of red-fluorescent cells was not observed (Fig. 8). Therefore, CrePA provides effective resistance against MGEs encoding Acr proteins that actively inhibit the ADP1 CRISPR-Cas, although the MGEs are actually not targeted by CRISPR-Cas.

**Fig. 8.**
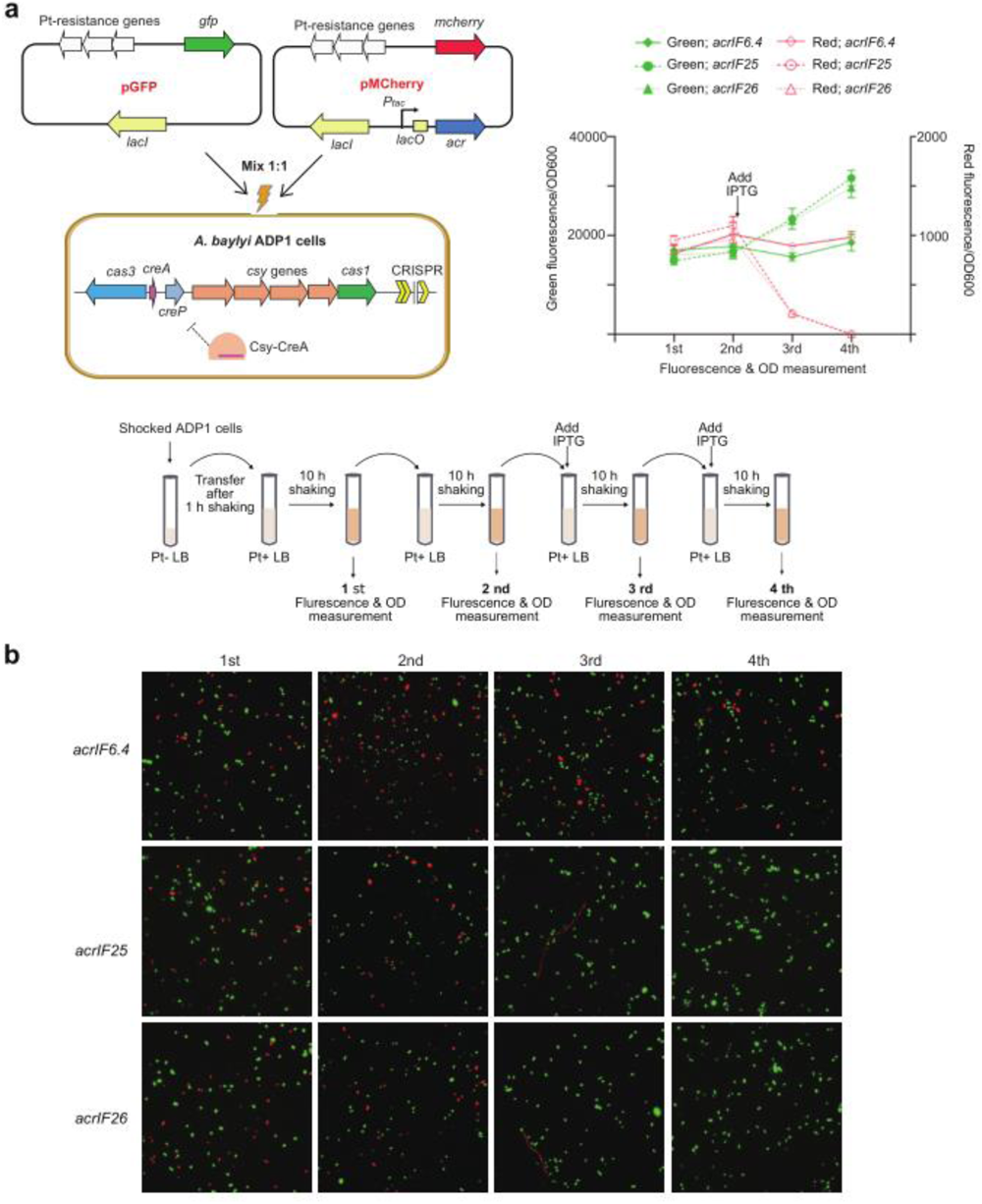
Stability of Acr-expressing plasmids in ADP1 cell population. **a.** Co-culturing of ADP1 cells containing an Acr-expressing plasmid (pMCherry) with those containing a plasmid not expressing Acr proteins (pGFP). The *acr* gene (*acrIF6.4*, *acrIF25*, or *acrIF26*) was controlled by an IPTG-inducible P*tac*. These two plasmids were mixed in a 1:1 ratio and introduced into ADP1 cells by electro-transformation. The shocked cells were initially grown in non-selective medium (with potassium tellurite, Pt-) to allow the expression of resistance genes for 1h, and then transferred to selective medium (Pt+) for cultivation. IPTG was added after the second sub-inoculation. The schedule of culturing and sampling is depicted. The plots illustrate the measurements of green fluorescence and red fluorescence in the collected samples. **b.** Fluorescence microscopy was employed to observe the ADP1 cells sampled from the co-culturing assay.

## DISCUSSION

In addition to its primary role in immunity, mounting evidence supports that CRISPR-Cas effectors have a secondary role in gene regulation. This regulatory role is usually guided by crRNA-like RNAs (crlRNAs) derived from standalone CRISPR repeats^30^. Several species of crlRNAs have been characterized, including CreR (Cas-regulating RNA, actually discovered during a systematic search for CreTA analogs) molecules from type I and V-A systems^31^ and tracr-L (long isoform of trans-activating CRISPR RNA) RNAs from type II-A systems^32^. These two major crlRNA species both mediate the autorepression of their corresponding CRISPR-Cas effector, whether multi-subunit or single-protein. Remarkably, this RNA-based Cas autoregulation reduces energy burden and the risk of autoimmunity while ensuring a required dynamic level of Cas proteins, which is critical for effective immunity when the cellular concentration of crRNAs changes or when Cas proteins are inhibited by some Acr proteins^31^. Therefore, these two crlRNA species play a role in mitigating the potential downsides of CRISPR-Cas during bacterial evolution.

CreA RNAs are a unique subset of crlRNAs that direct CRISPR-Cas effectors to transcriptionally suppress a deleterious toxin called CreT. Here, we demonstrated that CreTA possesses the potential as a broad-spectrum anti-anti-CRISPR mechanism. Acr proteins or Racr RNAs are frequently encoded and disseminated widely by various MGEs within the bacterial population. In a population of cells lacking CreTA, CRISPR-Cas is highly susceptible to the MGE-encoded Acr elements, and as a result, MGEs expressing Acr continuously diminish the proportion of cells with active CRISPR-Cas systems (Supplementary Fig. 8). While in the presence of CreTA, cells with MGEs expressing Acr elements that inhibit the CRISPR effector fail to suppress the expression of CreT (or CreP) toxins, resulting in cell death/dormancy or defects in cell division (Supplementary Fig. 8). These detrimental effects on the host cells can effectively eradicate Acr-encoding MGEs from the population, as observed in Fig. 8. Importantly, our data highlights the fact that a specific CreTA module (CrePA) can be activated by a wide variety of Acr proteins that inhibit the target DNA binding activity of CRISPR effectors (Fig. 5), thereby illustrating the broad-spectrum capacity of CreTA in resisting Acr elements. Leveraging this property of CreTA, we demonstrated the feasibility of screening for Acr proteins that effectively target a given CRISPR-Cas system (Fig. 6). The observation that Acr-encoding MGEs were expelled from the population of cells with a CreTA module (Fig. 8) indicates that such a CRISPR-regulated TA element provides host cells with population-level immunity against Acr-encoding MGEs, which interestingly are not targeted by any CRISPR spacers. In other words, the regulatory function of CRISPR-Cas confers herd immunity rather than targeted immunity in this scenario. Although the discovery of CreTA is limited to a subset of CRISPR-Cas systems, partly due to the challenges in predicting these highly diverse small genetic elements, it is highly probable that CRISPR-Cas has evolved in bacteria to regulate important physiological processes. For example, Cas9 has been found to transcriptionally regulate a regulon related to virulence in the pathogenic bacterium *Francisella novicida*, and this regulation relies on a crlRNA known as small CRISPR-associated RNA (scaRNA)^33^. Given these physiological benefits, the inhibition of CRISPR-Cas by Acr-encoding MGEs may similarly incur disfavored effects on host cells, leading to their extinction from the bacterial population after several generations. Therefore, it is tempting to speculate that the secondary gene regulation role of CRISPR-Cas could potentially confer herd immunity to host cells against Acr-encoding MGEs on a broader scale.

In addition, we discovered a unique CreTA module that utilizes a toxic protein (CreP) likely derived from phage proteins. This finding suggests that CRISPR-Cas has the trend to harness various toxic elements, whether they are proteinic or RNA-based, endogenous or derived from phages, to generate CreTA modules.

In summary, our data underlines the genetic diversity of CreTA elements and the versatility of the diverse crRNA-like regulatory RNAs. It is important to note that the anti-anti-CRISPR role of CRISPR-regulated TA modules, which acts on the population level, highlights the potential of Cas effectors in providing host cells with herd immunity against Acr-encoding MGEs, which is not dependent on CRISPR targeting.

## METHODS

### Bacterial strains and growth conditions

The strains used in this study are listed in Supplementary Data. 1*. E. coli* DH5α strain was utilized for plasmid construction, and *E. coli* BL21 (DE3) strain was used for protein expression and purification. *A. baylyi* ADP1, *A. baumannii* AYE, and *A. baumannii* ATCC 17978 cells were used to investigate the role of *A. baylyi* ADP1 *crePA* and the inhibitory effect of Acrs on CRISPR-Cas. All bacterial strains were cultured at 37°C in Lysogeny Broth (LB) media containing 10 g/L tryptone, 5 g/L yeast extract, 10 g/L NaCl. Solid plates were prepared using LB medium supplemented with 15 g/L agar, while liquid cultures were grown with agitation at 200 rpm. When necessary, the media were supplemented with uracil (50 μg/ml), 5-fluoroorotic acid (5-FOA; 50 μg/ml), isopropyl-b-D-thiogalactoside (IPTG, 0.5 mM), kanamycin (50 μg/ml), potassium tellurite (30 μg/ml), or apramycin sulfate (50 μg/ml).

### Plasmid construction and transformation

The plasmids, oligonucleotides, and synthetic genes used in this study are listed in Supplementary Data. 3. The double-stranded DNA fragments were amplified using Phanta Super-Fidelity DNA polymerase (Vazyme Biotech, Nanjing, China), followed by assembly into a pre-digested plasmid using the Trelief® Seamless Cloning Kit (Tsingke, Beijing, China) by the Gibson assembly strategy. Point mutations were introduced by overlap extension polymerase chain reaction (PCR), as previously described, and the engineered plasmids were verified by DNA sequencing.

Plasmids were transformed into *E. coli* DH5α, and *E. coli* BL21 (DE3) using commercially available competent cells. *A. baylyi* ADP1, *A. baumannii* AYE and *A. baumannii* ATCC 17978 cells were transformed using the electro-transformation method, unless otherwise specified. To prepare the electroporation-competent cells, a single colony was inoculated into liquid medium and the culture was grown overnight at 37 °C with agitation at 200 rpm. The overnight culture was then sub-inoculated into fresh LB medium with a ratio of 1:100 and cultured until the OD_600_ reached 0.6. The cultures were then transferred onto ice, centrifuged at 4000 rpm and washed three times with 10% glycerol. Then 300 ng of plasmids were transformed into 100 μl of competent cells using the bacteria program on the MicroPulser Electroporator (Bio-Rad, CA, USA). After 1 h recovery culture at 37°C in 500 μl of LB, the culture was serially diluted tenfold with LB and plated on selected plates containing potassium tellurite and/or other required antibiotics. Transformation efficiency was calculated by counting the colonies on the plates and multiplying the value by the dilution ratio. Each assay was conducted with three biological replicates. The mean and standard deviation of the transformation efficiency (colony forming units per μg of plasmid DNA, CFU/μg) were calculated from log-transformed data.

### Strain construction

The *A. baylyi* ADP1 mutant Δ*crePA* was generated using a *pyrF*-based efficient genetic manipulation platform, as previously described.^34^ The gene knockout plasmid was introduced by electro-transformation, and the single-crossover colonies were obtained by potassium tellurate screening. At least two PCR-validated colonies were inoculated onto the stationary phase with the addition of uracil and 5-FOA. Colonies were further screened and validated using replica plating and PCR amplification. Finally, DNA sequencing was performed to confirm the genotype.

### Searching for *crePA* homologs and analogs

To find *crePA* homologs, we searched the NCBI protein database using the ADP1 CreP and Csy1 proteins and retrieved the *cas3-csy1* intergenic regions (IGRs) that interfere with their most relevant homologs. By manually checking these IGR sequences, we identified 49 *crePA* modules (Supplementary Data. 1).

### Dilution plating assay

Cells were cultured overnight and sub-inoculated to reach the stationary phase after approximately 6 hours. For the experiments shown in Fig. 1c and Fig. 2c, the stationary culture was directly subjected to serial dilution and then plated on solid LB plates with or without 0.5 mM IPTG. In Supplementary Fig. 2C 0.5 mM IPTG or an equal volume of LB medium (as a control) was added onto the stationary cultures. After 0, 0.5, 1, 2, 4, or 6 hours, the cells were harvested, washed, and serially diluted with LB medium. Subsequently, 2 μl of each dilution were spotted onto solid LB plates with or without 0.5 mM IPTG.

### Cell morphology with scanning electron microscopy (SEM)

To prepare samples for SEM analysis, *A. baylyi* ADP1 cells containing either P*_tac_* promoter-induced *creP* expression or mutant promoter not expressing *creP* were cultured overnight. The cultures were sub-inoculated into LB medium supplemented with 1 mM IPTG and incubated at 37°C with shaking at 200 rpm for 12 h. After the incubation, the cells were harvested and washed twice. The final cell pellets were then pre-fixed with 3% glutaraldehyde (in 1X PBS) for 8 h at 4°C. To remove the excess glutaraldehyde, the cells were washed three times with 1X PBS. The cells were then dehydrated using a series of ethanol (50%, 70%, 85%, 95%, and 100%) for 15 min each. Finally, the samples were dried using the critical point drying method and then sputter-coated with gold. The samples were examined using a Hitachi SU8010 scanning electron microscope (Tokyo, Japan).

### RNA extraction and small RNA sequencing

*A. baylyi* ADP1 log phase cells were harvested for RNA extraction. Total RNA was extracted using the TRIzol reagent (Thermo Fisher Scientific, MA, USA) according to the manufacturer’s protocol. It was then purified using the phenol:chloroform method, followed by precipitation with an equal volume of ethanol. The purified RNA was stored at −80°C. For small RNA sequencing, 50 μg RNA was treated with T4 polynucleotide kinase (New England Biolabs, MA, USA) according to the manufacturer’s instructions at 37°C. The RNA was purified using the phenol-chloroform method and then precipitated with an equal volume of isopropanol and 0.1 volume of 3 M sodium acetate. The resulting RNA sample was redissolved in RNA-free water prior to sequencing. A small RNA library was constructed using the NEXTFLEX Small RNASeq Kit (Bioo Scientific, TX, USA), targeting RNA molecules between 30 and 300 nt. The library was then subjected to Illumina HiSeq sequencing (paired-end, 150 bp reads). The resulting raw data reads were adapter removed and were mapped to the *creA* sequence using previously reported Perl scripts.

### Primer extension analysis

Primer extension analysis for *A. baylyi* ADP1 was performed based on the previously described method^18^, using the 5’-FAM (6-carboxyfluorescein)-labeled *gfp*-specific primer as the reverse transcription primer. Briefly, 5 μg of the total RNA from *A. baylyi* ADP1 was first digested with RQ1 DNase (Promega, WI, USA), followed by reverse transcription to complementary DNA (cDNA) using 30 enzyme units(U) of the Moloney Murine Leukemia Virus reverse transcriptase (MMLV-RT) (Promega, WI, USA). The extension products were then analysed using the ABI3730xl DNA Analyzer (Thermo Fisher Scientific, MA, USA), and the results were viewed using Peak Scanner Software v1.0.

### Fluorescence measurement

*A. baylyi* ADP1 cells expressing *gfp* were grown until the late exponential phase, and the OD_600_ and fluorescence were measured simultaneously using the Synergy H4 Hybrid multimode microplate reader (BioTeck, VT, USA). At least three individual colonies were randomly selected and cultured for each experimental setting. The fluorescence/OD_600_ ratio was calculated for each of the three individual biological samples, and the mean and standard deviation were determined accordingly. Finally, a two-tailed Student’s t-test was performed to assess significance.

### Search for candidate *acr* genes from the *Acinetobacter* genomes

The search for *acr* genes was based on the fact that *acr* genes tend to occur in clusters and are located upstream of *aca* genes. First, we selected seven predicted *acr* genes from *Acinetobacter* (AcrIFs 6.2, 6.3, 6.4, 6.5, 11.2, 11.3, and 11.4) from the website http://guolab.whu.edu.cn/anti-CRISPRdb/ and examined their upstream and downstream sequences. We then discovered a conserved gene, *aca10*, located downstream of AcrIF 6.2, 11.3, and 11.5. To complement this finding, we performed a comprehensive search and analysis of *Acinetobacter* strains from NCBI, taking into account our custom *aca10* and the reported *aca7* gene. As a result, we discovered several potential *acr* genes located in close proximity to the *aca* genes. These genes were predicted as candidate *acr* genes using the website http://guolab.whu.edu.cn/anti-CRISPRdb/. Notably, AcrIFs 25 and 26 were among the predicted *acr* genes.

### Screening of Acr proteins with the CrePA module

To demonstrate the anti-anti-CRISPR function of *crePA,* the reported and predicted *acr* genes were synthesized and assembled to construct pACR. The pACR plasmid was then transformed into the host bacterial electroporation-competent cells expressing the Csy complex and *crePA*. For each experimental condition, three biological replicates (derived from different colonies) were included, and the transformation efficiency, mean, and standard deviation were determined. Finally, a two-tailed Student’s t-test was performed to determine significance. For the assay shown in Fig. 5, *A. baumannii* ATCC 17978(lacking CRISPR-Cas)was used as the host with an artificial plasmid pCAS-TA expressing the *P. aeruginosa* PA14 Csy complex and ΨR-replaced *crePA* as the host. For the assays shown in Supplementary Fig. 4, 6B, and 6C, *A. baylyi* ADP1 or Δ*crePA* was used as the host.

### Anti-CRISPR assay for Acr proteins

The CRISPR targeting assay was used to demonstrate the anti-CRISPR function of the Acrs in *A. baylyi* ADP1. First, plasmids containing different *acr* genes were individually transformed into *A. baylyi* ADP1. Subsequently, another plasmid containing the PAM sequence and spacer1 of *A. baylyi* ADP1 CRISPR, which served as the CRISPR target, was transformed into each strain to assess the immune function of CRISPR-Cas. A plasmid lacking the target site was used as a control. Three biological replicates (derived from different colonies) were included for each experimental condition, and the transformation efficiency, mean, and standard deviation were determined. Finally, a two-tailed Student’s t-test was performed to determine significance.

### Acrs abolish P*_creT_* inhibition

Fluorescence measurement was used to show that Acr proteins can affect the inhibition of P*_creT_* promoter by *creA* by interfering with the Csy-CreA complex. First, a plasmid expressing *gfp* under the P*_creT_* promoter was transformed into *A. baylyi* ADP1. Then, another plasmid capable of inducing acr gene expression was reintroduced into these strains. Individual colonies were randomly selected and cultured for 4 h. Subsequently, 0.5 mM IPTG was added to each culture. Fluorescence was measured after 6 hours of incubation.

### Protein expression and purification

Plasmids pET28a-csy, pET28a-acrIF25, and pET28a-acrIF26 were transformed into *E. coli* BL21 (DE3) to express the Csy complex, AcrIF25, and AcrIF26, respectively. *E. coli* BL21 was then inoculated into 1 L LB medium (50 μg/mL kanamycin) and cultured on a shaker at 37°C. Cultures were grown to an optical density (OD600 nm) of 0.75 and then induced with 0.5 mM IPTG for 8 h at 22°C (Csy complex) or 0.1 mM IPTG for 16 h at 22°C (AcrIF25/26). Protein purification was performed on a HisTrap HP column (Cytiva) at 4°C according to standard protocols. Harvested cells were collected by centrifugation (5000 rpm, 30 min, 4°C) and resuspended in 30 mL of binding buffer (20 mM HEPES pH = 7.5, 0.5 M NaCl, 20 mM imidazole, 5% glycerol). The cells were then sonicated on ice and after centrifuged (15000 g, 1 h, 4°C) to obtain the supernatant.

The supernatant was applied to HisTrap HP column equilibrated with the binding buffer and washed with the same buffer containing 20 mM imidazole. Proteins were eluted with the binding buffer containing 250 mM imidazole. The protein concentration was determined by the BCA method using bovine serum albumin (BSA) as a standard. The purified protein was then stored at −80°C.

### Tandem affinity chromatography (TAP)

The interaction of CreP with FtsZ and other protein partners was detected tandem affinity chromatography (TAP) using His-tag affinity chromatography coupled with co-immunoprecipitation (CoIP). For the His-tag affinity chromatography, approximately 20 mg of cell extract was purified on a His-Trap HP column according to the manufacturer’s procedure. The product was concentrated and imidazole removed using Amicon Ultrafra-30 concentrators (Millipore). The concentrated product was then immunoprecipitated using anti-Flag co-immunoprecipitation. For anti-Flag co-immunoprecipitation, prewashed anti-FLAG M2 magnetic beads (Sigma) were added to the concentrated His-Trap eluted product and incubated 8 h at 4°C with gentle shaking. The antigen-bound magnetic beads were washed five times with lysis buffer and eluted with 2.5 volumes of 3× FLAG peptide (150 ng/μl, Sigma) for 3 h at 4°C with gently shaking. The TAP products were then separated on a 12% SDS-PAGE gel and identified by Western blot using the anti-FtsZ antibody or by mass spectrometry (MS) according to the previous procedure.^35^

### Western blot

Western blot was performed to determine the presence of FtsZ in the TAP products of tagged CreP. A mouse polyclonal antiserum against FtsZ was raised by MBL International Corporation. The TAP products were separated on 12% SDS-PAGE and then transferred to a nitrocellulose membrane. The anti-FtsZ antisera (1: 5 000) were used first and then a horseradish peroxidase (HRP)-linked secondary conjugate at 1:5000 dilution was used for immunoreaction with the anti-FtsZ antiserum. Immune-active bands were visualized by an Amersham ECL Prime Western blot detection reagent (GE Healthcare).

### Size exclusion chromatography (SEC)

Affinity-purified proteins were fractionated by SEC using a GE Life Sciences Superdex 200 10/300 column. Fractions were collected in 0.5 ml volumes and monitored by optical density at 280 nm. SDS-PAGE gels were stained with silver nitrate or Coomassie Blue R250 for protein identification. For interaction experiments, purified Csy complex and AcrIF25/AcrIF26 were incubated together for 1 hour at 4°C before fractionation by SEC and co-eluting proteins were identified by SDS-PAGE. The protein mixture was then fractionated on the SEC column at 4°C. A fraction of the input (the purified Csy complex and AcrIF25/AcrIF26) was also retained for SDS-PAGE analysis.

### Electrophoretic mobility shift assay (EMSA)

To generate dsDNA, the biotin-labelled strand was heated to 98°C in the presence of a twofold excess of an unlabeled complementary strand and allowed to slowly return to room temperature. Specifically, non-target dsDNA without biotin labelling was generated by the same method described above, except that both strands were unlabeled. All full-length single-stranded DNA molecules were synthesized by GenScrip (Nanjing, China) and listed in Supplementary Data. 3.

Csy complex-DNA binding reactions were performed in binding buffer (20 mM HEPES, pH 7.5, 1 mM MgCl_2_, 150 mM NaCl and 5% glycerol) at 37 °C for 15 min. The concentrations of Csy complex, dsDNA and AcrIF25/AcrIF26 proteins are indicated in the fig. legends. After the appropriate incubation, the reactions were resolved on native 6% polyacrylamide TBE gels. The free dsDNA and shifted protein-DNA complexes in the gels were then transferred to a nylon membrane and cross-linked using GS Gene Linker UV Chamber (Bio-Rad Laboratories, Hercules, CA). The nylon membrane was incubated with 20 μg/mL Proteinase K (Ambion) at 55°C for 2 h, then chemiluminescence was detected using a Chemiluminescent Nucleic Acid Detection Module kit (Thermo Scientific) by exposure on a Tanon-5200 Multi instrument (Tanon Science & Technology Co. Ltd., Shanghai, China).

### Crystallization and X-ray data collection

AcrIF25 was concentrated to 15 mg/ml in 25 mM Tris-HCl pH 7.5, 150 mM NaCl and 2 mM DTT. Crystals were grown by the suspended droplet vapor diffusion method. Crystals of AcrIF25 were grown at 18 ℃ by mixing 0.5 ml of protein (15 mg/ml) with 0.5 ml of reservoir solution. Crystals were grown to full size in approximately 1 to 2 weeks. Crystals were obtained in 20% PEG3350 and 0.2 M sodium sulphate solution. Crystals were fished into cryoprotectant solution containing 20% glycerol and quickly transferred to liquid nitrogen for quick freezing. X-ray diffraction data were collected on the BL10U2 beamline at the Shanghai Synchrotron Radiation Facility (SSRF). Data processing, including indexing, integration and scaling, was performed using autoPROC XDS software. The initial model was solved using PHENIX and manually refined using COOT. Data collection and structure refinement statistics are summarized in Supplementary Data. 2, and all structure plots were generated using PyMOL software.

### Cross-linking experiments

Using disuccinimidyl suberate (DSS) as a cross-linking agent, 20 mg of protein was mixed with DSS, incubated on ice for 2 hours, then Tris-HCl (final concentration 2.5 mM) was added to stop the reaction and placed on ice for 15 minutes. The reaction samples were run on SDS-PAGE and visualized by Coomassie Blue staining.

### DNA competition experiments

To determine the enhanced binding ability of Csy complex to non-target DNA facilitated by AcrIF25 and AcrIF26, the DNA competition reactions were performed. In the absence or presence of AcrIF25 or AcrIF26, the DNA competition binding ability of the Csy complex to biotin-labelled target DNA was compared with that of the unlabeled non-target DNA. EMSA assays were performed as described above.

### Stability of Acr-expressing plasmids in the ADP1 cell population

300 ng plasmids expressing *gfp* and 300 ng plasmids expressing *acr* and *mcherry* were mixed and transformed into *A. baylyi* ADP1. After incubation in LB medium for 1 h, the culture was inoculated into fresh LB medium (containing 10 μg/ml potassium tellurite) with 1%, and fluorescence was measured after 10 h of incubation. This procedure was repeated four times, with the addition of 0.5 mM IPTG on the third and fourth occasions.

### Bioinformatic analysis

Multiple sequence alignment of CreP and its homologs was performed using MEGA7. Protein structures of CreP, KilA, Gp07, AcrIF25, and AcrIF26 were predicted using the AlphFold2 webserver. RNA secondary structures were predicted using the RNAfold webserver. Promoter elements were predicted using the BPROM program (Softberry tool).

### Statistics & reproducibility

All experiments in the present manuscript were repeated at least three times independently with similar results.

## REPORTING SUMMARY

Further information on research design is available in the Nature Portfolio Reporting Summary linked to this article.

## DATA AVAILABILITY

All relevant data are included in the paper and/or its supplementary information files. Source data are provided as a Source Data file. The accession number for the coordinate and structure factor of AcrIF25 is PDB 8X24. The raw data for the RNA-seq experiments in Fig. 4 were deposited to the National Center for Biotechnology Information (NCBI) with the BioProject accession number PRJNA1036288. All strains and plasmids are available from the corresponding author upon request; requests will be answered within 2 weeks. Source data are provided with this paper.

## ACKNOWLEDGEMENTS

This work was supported by the Science & Technology Fundamental Resources Investigation Program [2022FY101100], the National Natural Science Foundation of China [32150020, 82225028, 32270092, 32022003, 32370090, 32200057, and 82172287], the National Key Research and Development Program of China [2021YFC2301403], the Youth Innovation Promotion Association of CAS [2020090], and the China National Postdoctoral Program for Innovative Talents [BX20220331].

## AUTHOR CONTRIBUTIONS

M.L. and R.W. conceptualized this study. M.L., R.W., X.S., J.L., and S.O. designed the experiments, with valuable suggestions from C.H.. X.S., R.W., and Q.X. constructed the plasmids and mutant strains with the assistance of F.C., C.L., and H.Z.. X.S. and R.W. conducted dilution platting assay, SEM, the fluorescence measurement and bacteria transformation assays. X.S., R.W., and H.Z. performed RNA-seq and primer extension. R.W., X.S., and Q.X. performed the bioinformatic analyses of CrePA and Acrs. Z.L. and X.S. conducted protein purification, Western blot, SEC, TAP, and EMSA assays with the assistance of J.L. and Jf.L. X.S. conducted the DNA competition experiments and cell population assays. J.W. conducted crystallization and X-ray diffraction data collection of AcrIF25. Formal analysis of results was done by X.S., R.W., Z.L., Q.X., J. L., and S.O.. M.L., J.L., S.O., and R.W. analyzed the data and supervised the project. M.L. wrote the paper, which was edited and approved by all authors.

## REFERENCES

1. Barrangou, R., and Horvath, P. A decade of discovery: CRISPR functions and applications. Nat. Microbiol. 2, 17092. 10.1038/nmicrobiol.2017.92. (2017).

2. Wiedenheft, B., Sternberg, S.H., and Doudna, J.A.. RNA-guided genetic silencing systems in bacteria and archaea. Nature 482, 331-338. 10.1038/nature10886. (2012)

3. Hille, F., Richter, H., Wong, S.P., Bratovič, M., Ressel, S., and Charpentier, E. The biology of CRISPR-Cas: backward and forward. Cell 172, 1239–1259. 10.1016/j.cell.2017.11.032. (2018).

4. Nussenzweig, P.M., and Marraffini, L.A. Molecular mechanisms of CRISPR-Cas immunity in bacteria. Annu. Rev. Genet. 54, 93-120. 10.1146/annurev-genet-022120-112523. (2020).

5. Jackson, S.A., McKenzie, R.E., Fagerlund, R.D., Kieper, S.N., Fineran, P.C., and Brouns, S.J. CRISPR-Cas: adapting to change. Science 356, eaal5056. 10.1126/science.aal5056. (2017).

6. Makarova, K.S., Wolf, Y.I., Iranzo, J., Shmakov, S.A., Alkhnbashi, O.S., Brouns, S.J., Charpentier, E., Cheng, D., Haft, D.H., and Horvath, P. Evolutionary classification of CRISPR–Cas systems: a burst of class 2 and derived variants. Nat. Rev. Microbiol. 18, 67-83. 10.1038/s41579-019-0299-x. (2020).

7. Sternberg, S.H., Richter, H., Charpentier, E., and Qimron, U. Adaptation in CRISPR-Cas systems. Mol. Cell 61, 797-808. 10.1016/j.molcel.2016.01.030. (2016).

8. Semenova, E., Jore, M.M., Datsenko, K.A., Semenova, A., Westra, E.R., Wanner, B., Van Der Oost, J., Brouns, S.J., and Severinov, K. (2011). Interference by clustered regularly interspaced short palindromic repeat (CRISPR) RNA is governed by a seed sequence. Proc. Natl. Acad. Sci. USA 108, 10098–10103. 10.1073/pnas.110414410.

9. Wiedenheft, B., van Duijn, E., Bultema, J.B., Waghmare, S.P., Zhou, K., Barendregt, A., Westphal, W., Heck, A.J., Boekema, E.J., and Dickman, M.J. RNA-guided complex from a bacterial immune system enhances target recognition through seed sequence interactions. Proc. Natl. Acad. Sci. USA 108, 10092–10097. 10.1073/pnas.1102716108. (2011).

10. Bondy-Denomy, J., Garcia, B., Strum, S., Du, M., Rollins, M.F., Hidalgo-Reyes, Y., Wiedenheft, B., Maxwell, K.L., and Davidson, A.R. Multiple mechanisms for CRISPR–Cas inhibition by anti-CRISPR proteins. Nature 526, 136-139. 10.1038/nature15254. (2015).

11. Borges, A.L., Davidson, A.R., and Bondy-Denomy, J. The discovery, mechanisms, and evolutionary impact of anti-CRISPRs. Annu Rev Virol 4, 37–59. 10.1146/annurev-virology-101416-041616. (2017).

12. Jia, N., and Patel, D.J. Structure-based functional mechanisms and biotechnology applications of anti-CRISPR proteins. Nat. Rev. Mol. Cell Biol. 22, 563-579. 10.1038/s41580-021-00371-9. (2021).

13. Yin, P., Zhang, Y., Yang, L., and Feng, Y. Non-canonical inhibition strategies and structural basis of anti-CRISPR proteins targeting type I CRISPR-Cas systems. J. Mol. Biol. 435, 167996. 10.1016/j.jmb.2023.167996. (2023).

14. Camara-Wilpert, S., Mayo-Muñoz, D., Russel, J., Fagerlund, R.D., Madsen, J.S., Fineran, P.C., Sørensen, S.J., and Pinilla-Redondo, R. Bacteriophages suppress CRISPR–Cas immunity using RNA-based anti-CRISPRs. Nature 623, 601–607. 10.1038/s41586-023-06612-5. (2023).

15. Faure, G., Shmakov, S.A., Yan, W.X., Cheng, D.R., Scott, D.A., Peters, J.E., Makarova, K.S., and Koonin, E.V. CRISPR-Cas in mobile genetic elements: counter-defence and beyond. Nat. Rev. Microbiol. 17, 513–525. 10.1038/s41579-019-0204-7. (2019).

16. Li, M., Gong, L., Cheng, F., Yu, H., Zhao, D., Wang, R., Wang, T., Zhang, S., Zhou, J., and Shmakov, S.A. Toxin-antitoxin RNA pairs safeguard CRISPR-Cas systems. Science 372, eabe5601. 10.1126/science.abe5601. (2021).

17. Cheng, F., Wang, R., Yu, H., Liu, C., Yang, J., Xiang, H., and Li, M. Divergent degeneration of *creA* antitoxin genes from minimal CRISPRs and the convergent strategy of tRNA-sequestering CreT toxins. Nucleic Acids Res. 49, 10677–10688. 10.1093/nar/gkab821. (2021).

18. Wang, R., Shu, X., Zhao, H., Xue, Q., Liu, C., Wu, A., Cheng, F., Wang, L., Zhang, Y., and Feng, J. Associate toxin-antitoxin with CRISPR-Cas to kill multidrug-resistant pathogens. Nat. Commun. 14, 2078. 10.1038/s41467-023-37789-y. (2023).

19. Richter, C., Dy, R.L., McKenzie, R.E., Watson, B.N., Taylor, C., Chang, J.T., McNeil, M.B., Staals, R.H., and Fineran, P.C. Priming in the Type IF CRISPR-Cas system triggers strand-independent spacer acquisition, bi-directionally from the primed protospacer. Nucleic acids res. 42, 8516–8526. 10.1093/nar/gku527. (2014).

20. Iyer, L.M., Koonin, E.V., and Aravind, L. Extensive domain shuffling in transcription regulators of DNA viruses and implications for the origin of fungal APSES transcription factors. Genome Biol. 3, 1-11. 10.1186/gb-2002-3-3-research0012. (2002).

21. Hansen, E.B. Structure and regulation of the lytic replicon of phage P1. J. Mol. Biol. 207, 135–149. 10.1016/0022-2836(89)90445-2. (1989).

22. Das, A., Biswas, S., and Biswas, M. Expression of Phi11 Gp07 causes filamentation in *Escherichia coli*. The open Microbiol. J. 12, 107. 10.2174/1874285801812010107. (2018).

23. Cech, G.M., Szalewska-Pałasz, A., Potrykus, K., and Kloska, A. Virus–host interaction gets curiouser and curiouser. Part II: Functional transcriptomics of the E. coli DksA-deficient cell upon phage P1 *vir* infection. International journal of molecular sciences **22**, 6159. 10.3390/ijms22116159. (2021).

24. de Boer, P.A.J. Advances in understanding E. coli cell fission. Curr. Opin. Microbiol. 13, 730–737. 10.1016/j.mib.2010.09.015. (2010).

25. Cheng, F., Wu, A., Liu, C., Cao, X., Wang, R., Shu, X., Wang, L., Zhang, Y., Xiang, H., and Li, M. The toxin–antitoxin RNA guards of CRISPR-Cas evolved high specificity through repeat degeneration. Nucleic Acids Res. 50, 9442–9452. 10.1093/nar/gkac712. (2022).

26. Xie, Y., Zhang, L., Gao, Z., Yin, P., Wang, H., Li, H., Chen, Z., Zhang, Y., Yang, M., and Feng, Y. AcrIF5 specifically targets DNA-bound CRISPR-Cas surveillance complex for inhibition. Nat. Chem. Biol. 18, 670-677. 10.1038/s41589-022-00995-8. (2022).

27. Gao, Z., Zhang, L., Ge, Z., Wang, H., Yue, Y., Jiang, Z., Wang, X., Xu, C., Zhang, Y., and Yang, M. Anti-CRISPR protein AcrIF4 inhibits the type IF CRISPR-Cas surveillance complex by blocking nuclease recruitment and DNA cleavage. J. Biol. Chem. 298. 10.1016/j.jbc.2022.102575. (2022).

28. Yang, L., Zhang, L., Yin, P., Ding, H., Xiao, Y., Zeng, J., Wang, W., Zhou, H., Wang, Q., and Zhang, Y. Insights into the inhibition of type IF CRISPR-Cas system by a multifunctional anti-CRISPR protein AcrIF24. Nat. Commun. 13, 1931. 10.1038/s41467-022-29581-1. (2022).

29. Mukherjee, I.A., Gabel, C., Noinaj, N., Bondy-Denomy, J., and Chang, L. Structural basis of AcrIF24 as an anti-CRISPR protein and transcriptional suppressor. Nat. Chem. Biol. 18, 1417–1424. 10.1038/s41589-022-01137-w. (2022).

30. Shmakov, S.A., Barth, Z.K., Makarova, K.S., Wolf, Y.I., Brover, V., Peters, J.E., and Koonin, E.V. Widespread CRISPR-derived RNA regulatory elements in CRISPR-Cas systems. Nucleic Acids Res., gkad495. 10.1093/nar/gkad495. (2023).

31. Liu, C., Wang, R., Li, J., Cheng, F., Shu, X., Zhao, H., Xue, Q., Yu, H., Wu, A., and Wang, L. Widespread RNA-based *cas* regulation monitors crRNA abundance and anti-CRISPR proteins. Cell Host & Microbe 31, 1481–1493. e1486. 10.1016/j.chom.2023.08.005. (2023).

32. Workman, R., Pammi, T., Nguyen, B., Graeff, L., Smith, E., Sebald, S., Stoltzfus, M., Euler, C., and Modell, J. A natural single-guide RNA repurposes Cas9 to autoregulate CRISPR-Cas expression. Cell 184, 675-688. e619. 10.1016/j.cell.2020.12.017. (2021).

33. Ratner, H.K., Escalera-Maurer, A., Le Rhun, A., Jaggavarapu, S., Wozniak, J.E., Crispell, E.K., Charpentier, E., and Weiss, D.S. Catalytically Active Cas9 Mediates Transcriptional Interference to Facilitate Bacterial Virulence. Mol. Cell 75, 498-510. e495. 10.1016/j.molcel.2019.05.029. (2019).

34. Wu, S., Xu, R., Su, M., Gao, C., Liu, Y., Chen, Y., Luan, G., Xu, J., Wang, R. A pyrF-Based Efficient Genetic Manipulation Platform in Acinetobacter baumannii To Explore the Vital DNA Components of Adaptive Immunity for IF CRISPR-Cas. Microbiol. Spectr. 10(5), e01957–22. 10.1128/spectrum.01957-22. (2022).

35. Jia, J., Li, J., Qi, L., Li, L., Yue, L. and Dong, X. Post-transcriptional regulation is involved in the cold-active methanol-based methanogenic pathway of a psychrophilic methanogen. Environ. Microbiol. 23: 3773–3788. 10.1111/1462-2920.15420. (2021).

