## Supplementary information for "CRISPR-repressed toxin-antitoxin provides population-level immunity against diverse anti-CRISPR elements"

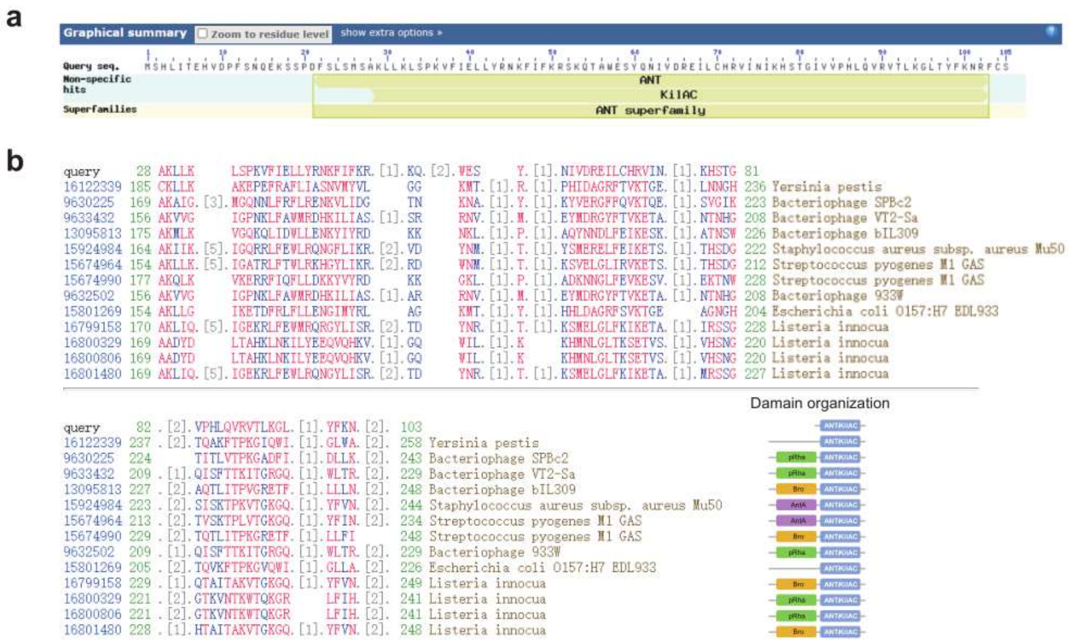


**Supplementary Fig. 1. CreP has a conserved KilAC domain that is likely derived from phage proteins**. **a**. The amino acid sequence and domain architecture of CreP protein. **b.** Multiple sequence alignment of representative proteins containing the KilAC domain (C-terminal domain of phage P1 KilA, COG3645). The query sequence is CreP, and the 13 homologous proteins are all from phages or prophages residing within bacterial genomes. The domain organization of each protein is depicted. ANTKilAC, the phage antirepressor protein KilAC domain; pRha, the phage regulatory protein RHA (COG3646); Bro, N-terminal domain of baculovirus BRO proteins (pfam02498); AntA, phage antirepressor protein (COG3561).

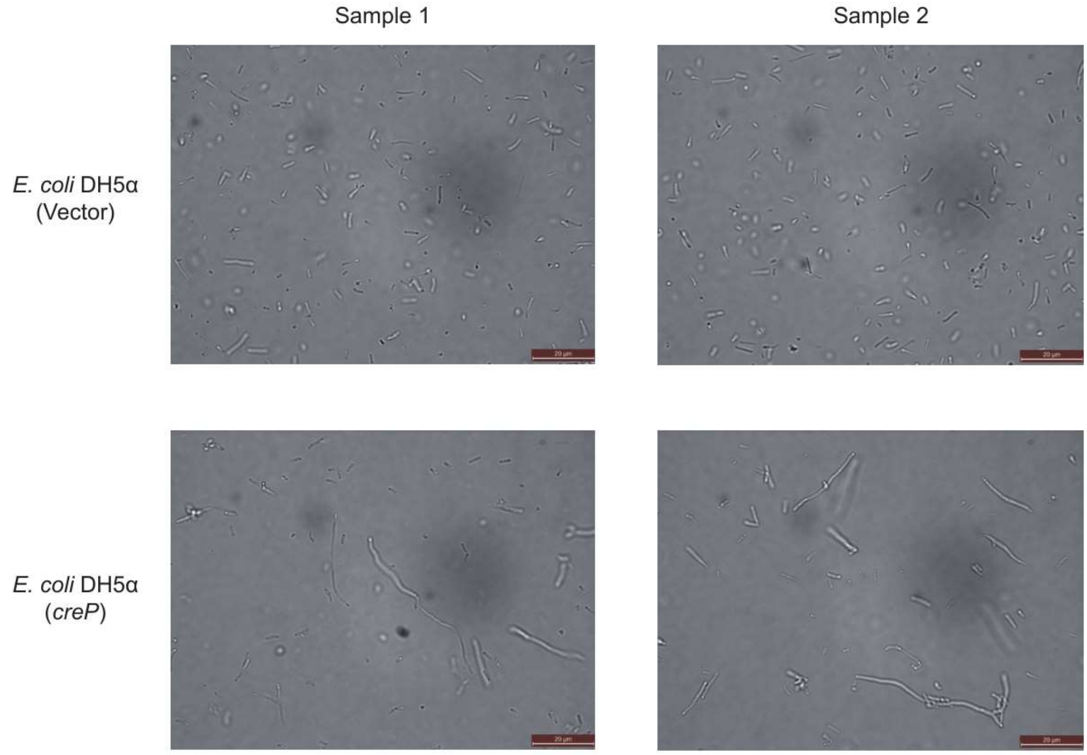


**Supplementary Fig. 2. Microscopy of *E. coli* cells expressing CreP.** *E. coli* DH5α cells over-expressing CreP or containing an empty vector were observed using an optical microscope. Scale bars, 20 µm.

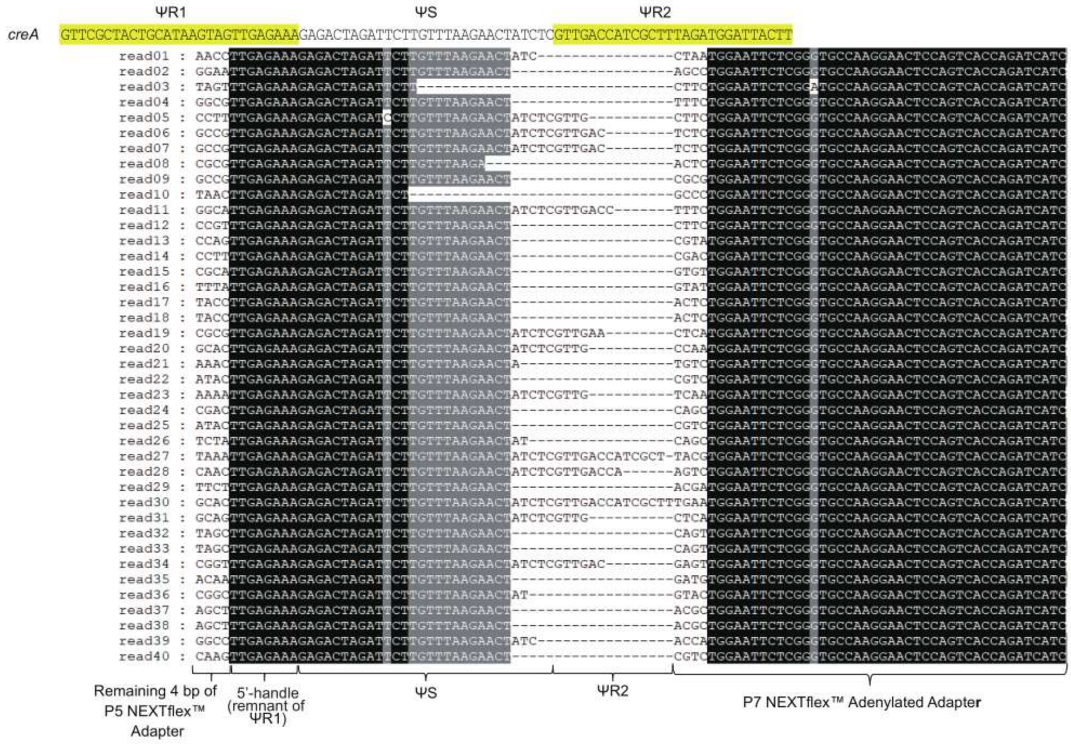


**Supplementary Fig. 3. Example reads of mature CreA RNA.** The sequences of 40 example reads were mapped to the DNA sequence of *creA* gene to determine their exact termini. The nucleotides corresponding to the adapters used during library construction are indicated.

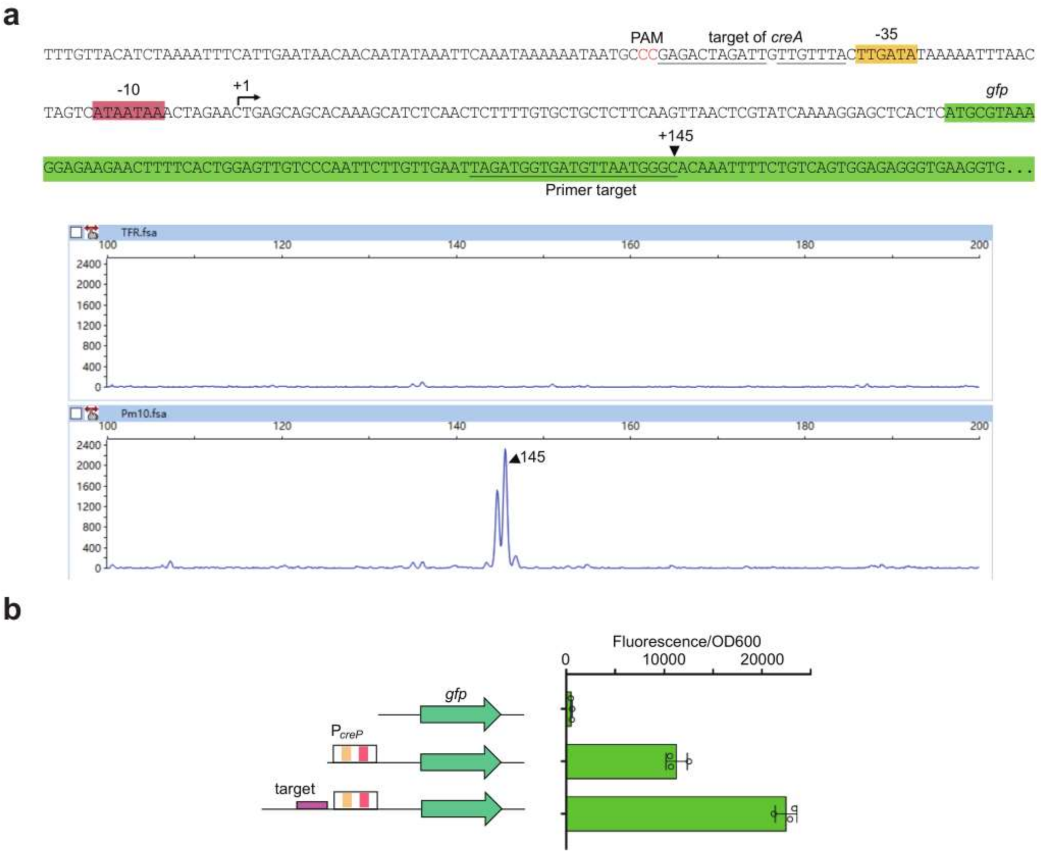


**Supplementary Fig. 4. Characterization of the *creP* promoter and its transcription start site (TSS). a.** Primer extension assay to determine the TSS of $P_{creP}$. $P_{creP}$ was first linked to a *gfp* gene and then introduced into *A. baylyi* ADP1 cells, of which total RNA was extracted for primer extension. The primer was designed against the *gfp* RNA transcript and 5'-labeled by FAM (see the Methods part). The cDNA products of primer extension assay were subjected to fragment size analysis. **b.** Assessing the strength of $P_{creP}$ with or without the upstream target site of CreA. Error bars, mean ± s.d. (n=3).

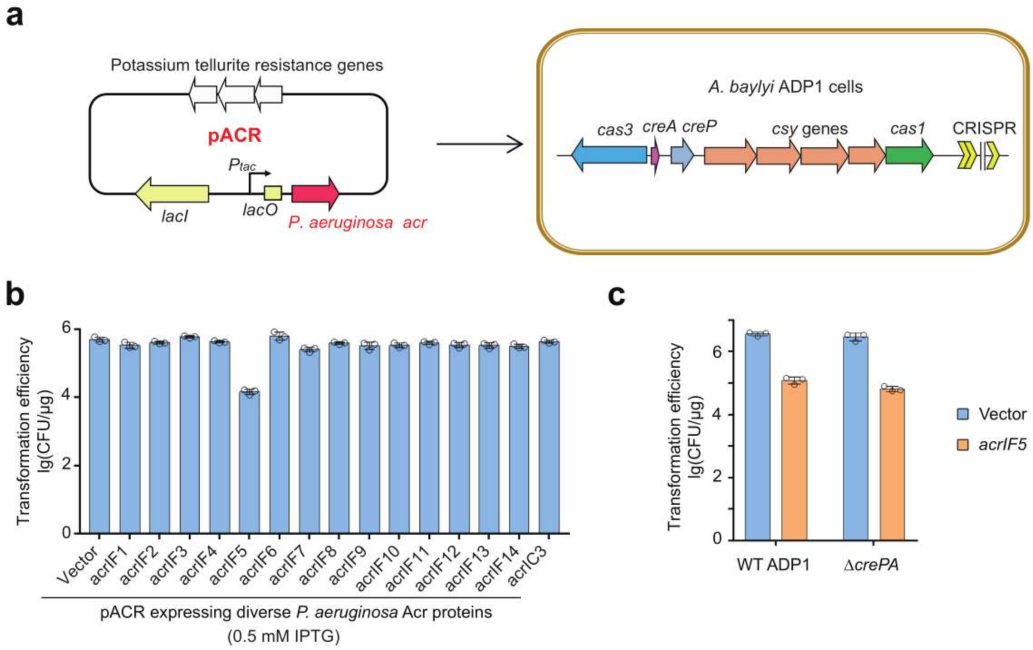


**Supplementary Fig. 5. The *crePT* module cannot be triggered by *P. aeruginosa* AcrIF proteins. a.** Schematic illustration of the assay. To test whether the toxicity of CrePA can be triggered by *P. aeruginosa* AcrIF proteins, plasmids expressing one of these proteins (controlled by an IPTG-inducible *tac* promoter) were introduced into wild-type *A. baylyi* ADP1 cells (containing *crePA*) by transformation. **b.** Transformation efficiency of ADP1 cells on plates containing 0.5 mM IPTG (inducing Acr expression). **c.** Transformation of wild-type ADP1 or Δ*crePA* cells by the plasmid expressing AcrIF5. Error bars, mean ± s.d. (n=3).

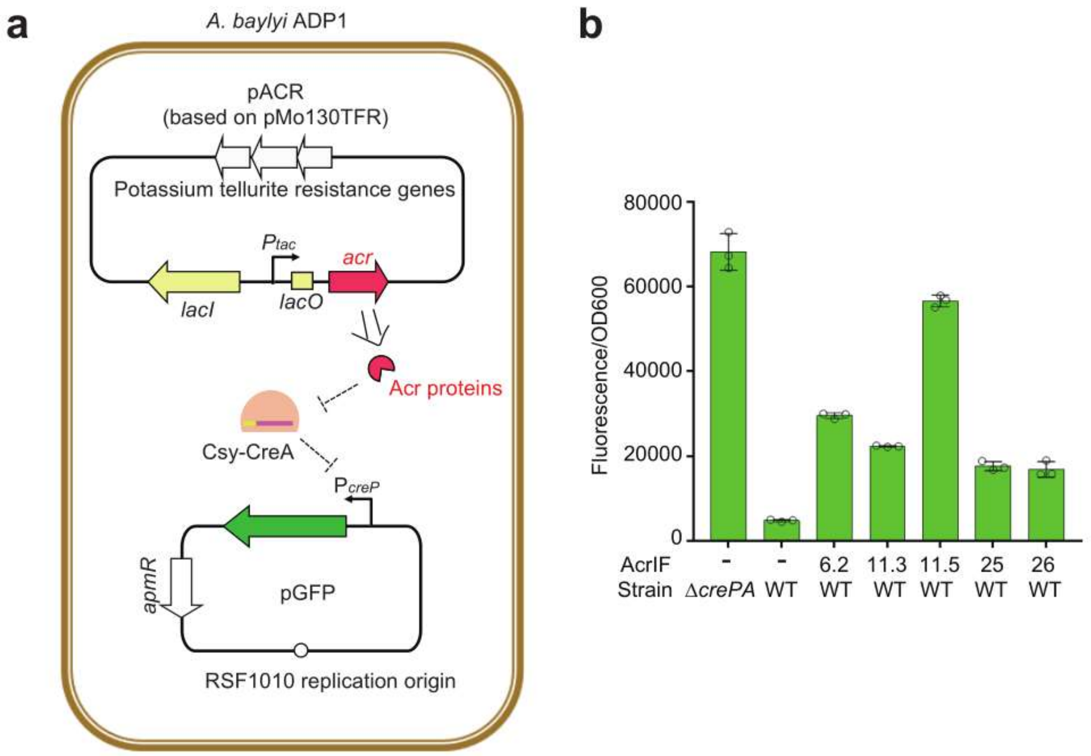


**Supplementary Fig. 6. Assessing the derepression effect of newly identified Acr proteins on P*creP*. a.** Schematic illustration of this assay. The assay involves two plasmids: pGFP expresses GFP under the control of $P_{creP}$ that is repressed by the Csy-CreA complex, while pACR expresses an Acr protein to inhibit the Csy complex, thus de-repressing $P_{creP}$. **b.** Measurement of fluorescence. The empty vector of pACR was included as a control (-). The Δ*crePA* control was used to show the strength of fully de-repressed $P_{creP}$ for comparison. Error bars, mean ± s.d. (n=3).

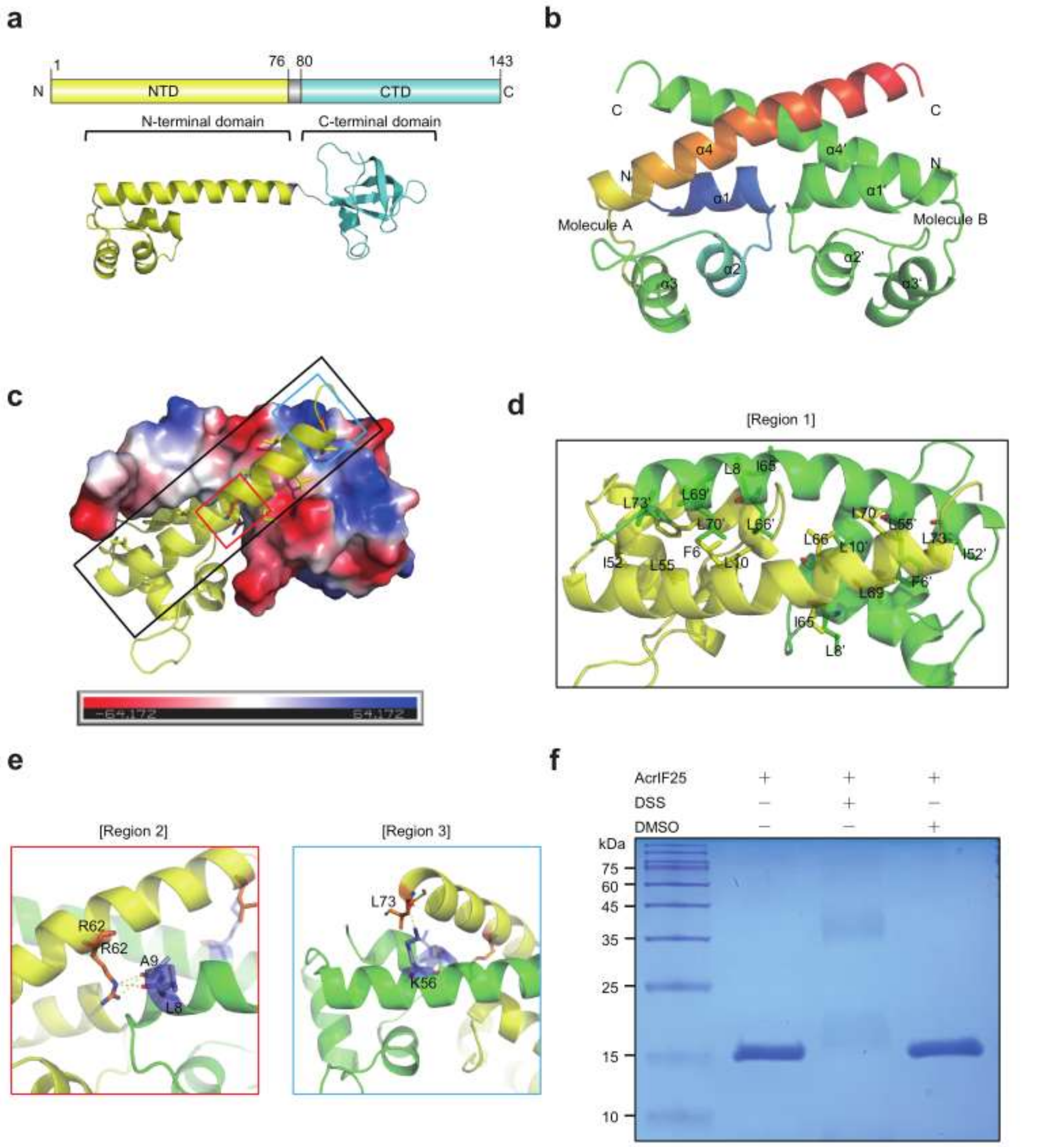


**Supplementary Fig. 7. Overall structure and polymerization of AcrIF25. a**. Structure of AcrIF25 predicted by AlphaFold2. A schematic diagram of the domain organization is also given. **b.** The cartoon structure of the dimer AcrIF25(1-76) is represented as a crystalline asymmetric unit. The monomer on the left (molecule A) has a chain color that gradually changes from blue to red in the spectrum from the N-terminus to the C-terminus, with secondary structures α1 - α4, respectively. The monomer on the right (molecule B) has a green color, with secondary structures α1' - α4', respectively. **c.** The initial dimer structure of AcrIF25(1-76) was generated and analyzed. Black boxes are used to indicate region 1, red boxes are used to indicate region 2, and blue boxes are used to indicate region 3. Three different close-up views of region 1 (**d**), region 2 (**e**), and region 3 (**e**) in the AcrIF25(1-76) dimer structure. Yellow dashed lines indicate hydrogen bonds. Yellow and orange sticks represent residues of molecule A, green and blue represent residues of molecule B. **f.** Polymerization state analysis of AcrIF25 by SDS-PAGE. The experimental group was incubated with AcrIF25 and DSS, one control group was incubated with AcrIF25 and the other control group was incubated with AcrIF25 and DMSO. The dimer state of AcrIF25 are framed in red.

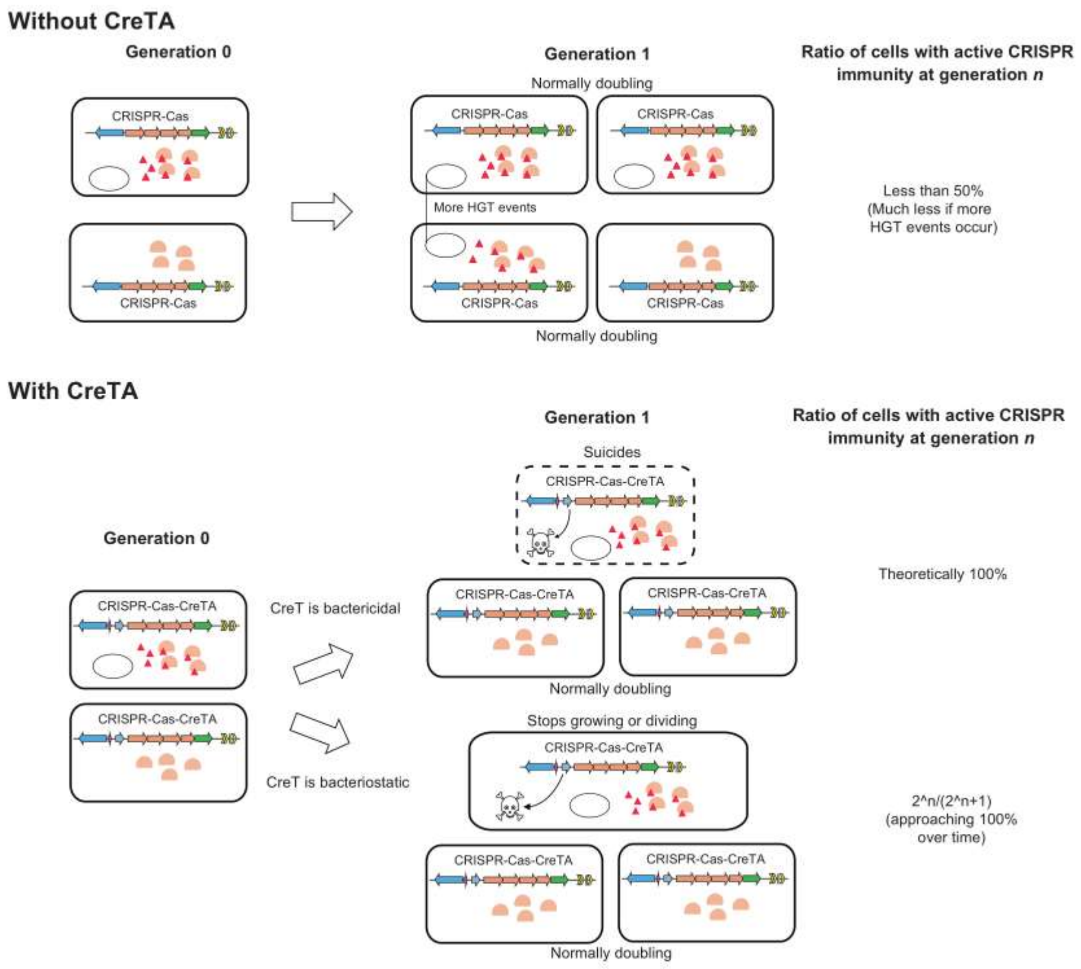


**Supplementary Fig. 8. The model of CreTA-mediated exclusion of *acr* genes from cell population.** In the absence of CreTA, the *acr* genes carried by MGEs can freely spread within the population, leading to a continuous decrease in the percentage of cells with active CRISPR immunity. However, in the presence of CreTA, cells that harbor MGEs expressing Acr proteins or Racr RNAs are unable to suppress the expression of CreT (or CreP) toxins, as the Acr elements inactivate the CRISPR effectors. As a result, these cells experience cell death, dormancy, or defects in cell division. These deleterious effects effectively exclude Acr-encoding MGEs from the population, thereby promoting the stable persistence of active CRISPR immunity in the bacterial population.
